# Exploratory neuroimmune profiling identifies CNS-specific alterations in COVID-19 patients with neurological involvement

**DOI:** 10.1101/2020.09.11.293464

**Authors:** Eric Song, Christopher M. Bartley, Ryan D. Chow, Thomas T. Ngo, Ruoyi Jiang, Colin R. Zamecnik, Ravi Dandekar, Rita P. Loudermilk, Yile Dai, Feimei Liu, Isobel A. Hawes, Bonny D. Alvarenga, Trung Huynh, Lindsay McAlpine, Nur-Taz Rahman, Bertie Geng, Jennifer Chiarella, Benjamin Goldman-Israelow, Chantal B.F. Vogels, Nathan D. Grubaugh, Arnau Casanovas-Massana, Brett S. Phinney, Michelle Salemi, Jessa Alexander, Juan A. Gallego, Todd Lencz, Hannah Walsh, Carolina Lucas, Jon Klein, Tianyang Mao, Jieun Oh, Aaron Ring, Serena Spudich, Albert I. Ko, Steven H. Kleinstein, Joseph L. DeRisi, Akiko Iwasaki, Samuel J. Pleasure, Michael R. Wilson, Shelli F. Farhadian

## Abstract

One third of COVID-19 patients develop significant neurological symptoms, yet SARS-CoV-2 is rarely detected in central nervous system (CNS) tissue, suggesting a potential role for parainfectious processes, including neuroimmune responses. We therefore examined immune parameters in cerebrospinal fluid (CSF) and blood samples from a cohort of patients with COVID-19 and significant neurological complications. We found divergent immunological responses in the CNS compartment, including increased levels of IL-12 and IL-12-associated innate and adaptive immune cell activation. Moreover, we found increased proportions of B cells in the CSF relative to the periphery and evidence of clonal expansion of CSF B cells, suggesting a divergent intrathecal humoral response to SARS-CoV-2. Indeed, all COVID-19 cases examined had anti-SARS-CoV-2 IgG antibodies in the CSF whose target epitopes diverged from serum antibodies. We directly examined whether CSF resident antibodies target self-antigens and found a significant burden of CNS autoimmunity, with the CSF from most patients recognizing neural self-antigens. Finally, we produced a panel of monoclonal antibodies from patients’ CSF and show that these target both anti-viral and anti-neural antigens—including one mAb specific for the spike protein that also recognizes neural tissue. This exploratory immune survey reveals evidence of a compartmentalized and self-reactive immune response in the CNS meriting a more systematic evaluation of neurologically impaired COVID-19 patients.

**One Sentence Summary:** A subset of COVID-19 patients with neurologic impairment show cerebrospinal fluid-specific immune alterations that point to both neuroinvasion and anti-neural autoimmunity as potential causes of impairment.

## Introduction

The causative pathogen of pandemic coronavirus disease 2019 (COVID-19), SARS-CoV-2, primarily causes a respiratory illness. Yet, in some patients, SARS-CoV-2 infection associates with severe and debilitating extrapulmonary symptoms, including neurological dysfunction(*1*). Indeed, about a third of patients with moderate to severe COVID-19 develop neurologic symptoms including anosmia, dysgeusia, headache, impaired consciousness, and seizure—only some of which is explained by systemic complications including hypercoagulability(*2*). While studies of cerebrospinal fluid (CSF) in affected patients only rarely detect SARS-CoV-2 RNA(*3–6*), the largest post-mortem neuropathological study to date found evidence for SARS-CoV-2 proteins in the brains of half of patients. However, there was little evidence of associated tissue damage(*7*). Collectively, these observations suggest mechanisms other than viral replication may contribute to neuropathology, and point to the need for a broad characterization of the neuroimmune milieu during the course of neurologic impairment of COVID-19 patients. In this exploratory study, we compared the innate and adaptive arms of the intrathecal and peripheral immune responses in patients with COVID-19 complicated by diverse neurological symptoms.

## Results

### Overview of study

Hospitalized COVID-19 patients with diverse neurological symptoms who underwent clinically-indicated lumbar puncture were consented for collection of surplus CSF to be used for research. Six participants with acute COVID-19, based on positive nasopharyngeal swab SARS-CoV-2 RT-qPCR, were enrolled (Cases 1 – 6, Supplementary Table 1). Neurological symptoms included encephalopathy (n=3), intractable headache (n=3), and seizure (n=1). All participants donated paired blood and CSF, except for one patient who did not donate blood. Lumbar punctures were performed on median hospital day 12.5 (range 2 - 43 days). Healthy controls were previously enrolled through a neuroinfectious disease biorepository at Yale prior to the COVID-19 pandemic, with CSF supernatant biobanked and frozen; these controls were included based on age and sex match to the COVID-19 cases (n=3). Because fresh CSF is required for single cell analyses, we also recruited additional uninfected control participants during the COVID-19 pandemic (n=3): 2 were healthy community-dwelling adults, and one was hospitalized for the work-up of frequent falls. These controls tested negative for SARS-CoV-2 by nasopharyngeal swab RT-PCR. Fresh CSF and blood samples were processed into cells from the CSF, CSF supernatant, peripheral blood mononuclear cells (PBMCs), and plasma (Figure 1a). CSF (n=5) and plasma (n=6) samples from COVID-19 cases were negative for SARS-CoV-2 RNA by RT-qPCR using the CDC primer-probe sets(*8*).

**Figure 1:**
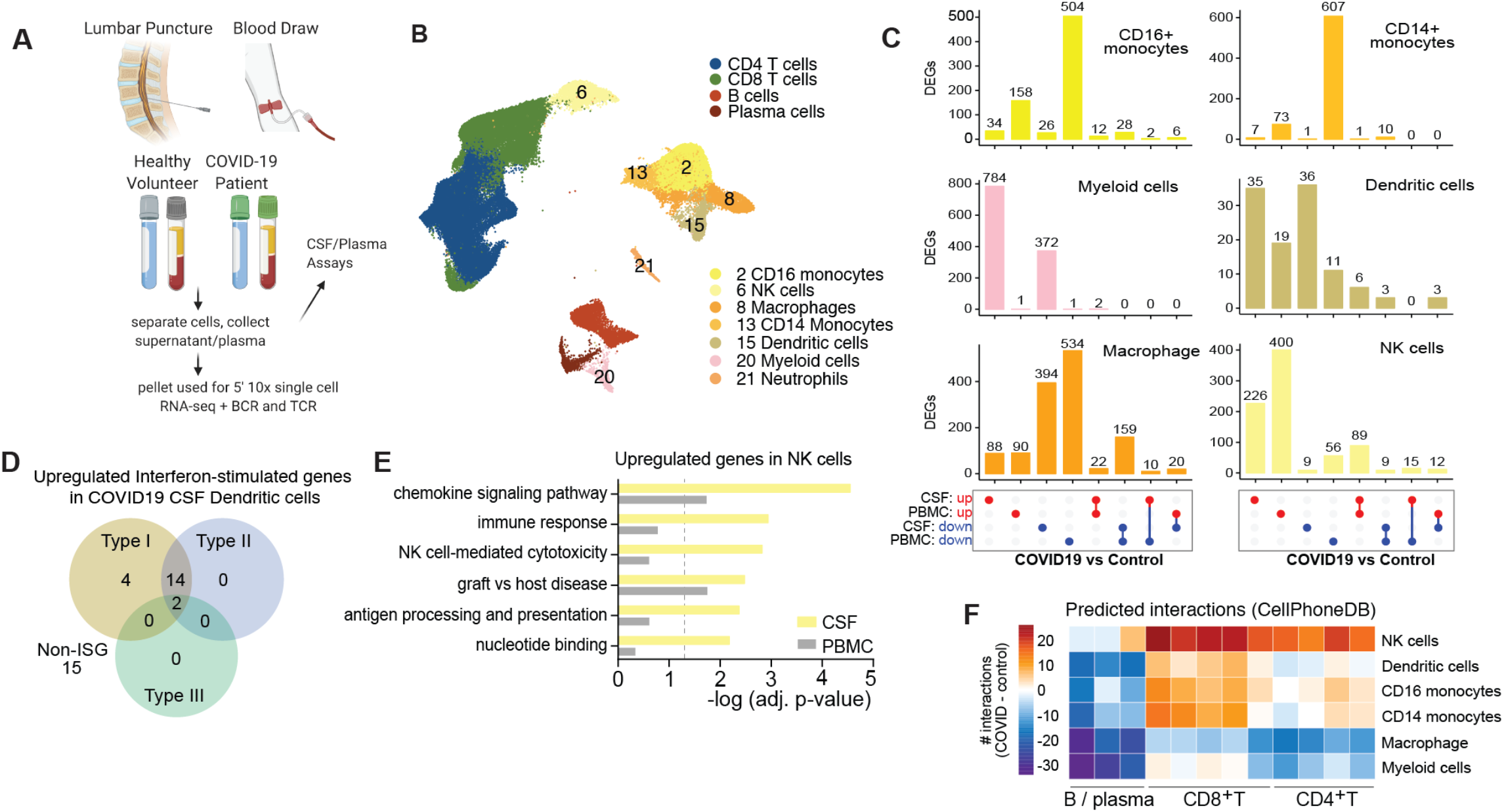
Distinct immunological landscape of CSF and PBMC in COVID-19 patients with neurological symptoms. (**A**) Schematic of study design. CSF and blood were collected from COVID-19 cases and healthy controls. PBMC and CSF cells were isolated, along with the CSF supernatant and plasma for downstream analysis. (**B**) UMAP projection of 10x single cell RNA-sequencing of CSF and PBMC of COVID-19 cases and healthy controls. Single cell transcriptomes were derived from a total of 42,495 PBMCs and 33,978 CSF cells from 5 COVID-19 cases and 11 controls, including 8 from Gate et al(*9*). (**C**) UpSet plot showing differentially expressed genes (DEGs) in innate immune cells of COVID-19 cases versus controls. The legend for each column is indicated in the bottom panel. Each column denotes the number of genes that are significantly (adjusted p value <0.05) up (red) or down (blue) regulated in COVID-19 tissue when compared to control. Connecting dots indicate the intersection of DEGs in CSF and PBMC. (**D**) Venn diagram depicting upregulated interferon-stimulated genes (ISG) and non-ISGs in dendritic cells in COVID-19 CSF compared to healthy control CSF based on Interferome database(*42*). (**E**) Gene ontology enrichment of genes upregulated in NK cells of COVID-19 patients in the CSF and peripheral blood. (**F**) Heat map depicting cell–cell interactions between innate immune cells and adaptive immune cells by clustering shown in Figure 1b. The difference in interaction strength (COVID-19 interaction minus control interaction) is color coded, and derived from log-scaled interaction counts using the CellphoneDB repository of ligands, receptors and their interactions(*10*).

### Transcriptional analysis reveals a coordinated innate immune cell response to COVID-19 in the CNS

To investigate the effect of SARS-CoV-2 infection on host immune cell gene expression, we performed single cell RNA sequencing with the 10X Genomics platform on 76,473 immune cells from the CSF and blood of patients hospitalized with acute COVID-19 (n = 5) and uninfected controls (n = 3 recruited in March – May 2020, and n = 8 healthy controls from Gate et al(*9*)). To test for the presence of intracellular virus, open reading frames of SARS-CoV-2 (spike, ORF3a, envelope, membrane glycoprotein, ORF6, ORF7a, ORF8, nucleocapsid, and ORF10) were added to the reference genome before alignment with CellRanger. No viral transcripts were detected in any of the CSF cells or PBMCs. We performed unsupervised cluster analysis and represented gene expression data from cases and controls in UMAP space. We identified distinct T cell, B cell, and myeloid cell populations (Figure 1b), characterized by gene expression patterns of canonical immune cell genes (fig. S1a). As expected, in both COVID-19 cases and controls B cells and neutrophils were enriched in the PBMC compartment compared to the CSF, whereas CSF was enriched for a population of myeloid cells rarely present amongst PBMCs (fig. S1b-c). To study CSF-specific gene expression patterns of immune cell populations in COVID-19 patients, we compared gene expression in CSF from COVID-19 cases and controls, and compared these differentially expressed genes to genes that were differentially expressed in the PBMCs between COVID-19 cases and controls. In this way, we found COVID-19 associated transcriptional changes that were unique to the CSF.

While CD16+ and CD14+ monocytes included the most differentially downregulated genes in the PBMCs of COVID-19 patients, dendritic cells, and CSF-predominant myeloid cells yielded the most differentially expressed genes in the CSF compartment (Figure 1c). While NK cells in the CSF and the peripheral blood demonstrated comparable changes in the number of differentially expressed genes in COVID-19 cases compared to controls, the genes affected were mostly unique to each compartment (Figure 1c, fig. S1d).

Using single cell differential gene expression analysis, we found evidence for activation of dendritic cells in the CSF of COVID-19 patients: 57% and 47% of the upregulated genes in these cells were classified as type 1 and type 2 interferon stimulated genes, respectively (Figure 1d). NK cells in the CSF of COVID-19 patients also specifically upregulated genes associated with NK cell activation (Figure 1e). Using CellphoneDB signaling network analysis(*10, 11*) we found that in the CSF of COVID-19 patients, dendritic and NK cells are predicted to have increased interaction with CD8+ and CD4+ T cells compared to what is predicted in the CSF of healthy patients (Figure 1f). We asked whether the predicted interactions between activated innate and adaptive immune cells in the CNS could be interpreted as driving the transcriptional changes observed in the CSF T cells of COVID-19 patients. Using NicheNet(*11*), we designated the CSF innate immune cells as “sender” cells and CD4+ T cells, CD8+ T cells and B cells as “receiver” cells, and found enrichment in CD4+ and CD8+ T cells of potential regulatory genes affecting chemokine receptors, integrins, and activation of immune cells, consistent with results from CellphoneDB analysis (fig. S2).

### T cells in the CSF display increased cellular activation during COVID-19

Because T cells were predicted to be the main recipients of innate-adaptive cross talk, we first subsetted and re-clustered CSF and blood T cells for further analysis (Figure 2a, fig. S3a-d). While the relative proportions of CSF T cell populations were generally conserved in COVID-19 patients compared to controls, among the PBMCs there was a decrease in the frequency of naïve CD4+ T cells (mean COVID-19 9.65%, healthy 18.219%; p =0.001) and an increase in effector CD8+ T cells (mean COVID-19 30.9%, healthy 16.245%; p= 0.02) (Figure 2b). However, while relative T cell populations were conserved in CSF of COVID-19 cases compared to control CSF, we found significant COVID-19 associated transcriptional changes in CSF T cells (fig. S3d). Effector CD4+ T cells in the CSF of COVID-19 patients showed significant transcriptional changes compared to CD4+ T cells in the PBMCs of COVID-19 cases. After excluding any genes that were also differentially expressed between T cells in the CSF and periphery of healthy individuals, we identified genes that were upregulated in both Th1 and Th2 CD4+ T cells from COVID-19 CSF (Figure 2d) and found enrichment for several gene pathways important for T cell activation, as well as engagement of IL-1 and IL-12 mediated signaling pathways (Figure 2e). Effector CD8+ T cells in the CSF were similarly enriched for genes involved with canonical immune response pathways when compared to CSF from healthy donors, including: 1) increased motility and cell adhesion, 2) differentiation/proliferation, and 3) effector programming (responses to IL-12, IL-1, IFN-g, and T cell co-stimulation), indicating the presence of a coordinated T cell based immune response in the CNS (Figure 2f-g). Overall, COVID-19 associated transcriptional changes were observed in both CD4+ and CD8+ T cells in the CSF, were agnostic to the effector state of the cells, and predicted cell-cell interactions that were unique to the CSF in COVID-19 cases, including T cell co-stimulation factors and trafficking interactions (fig. S4a-b).

**Figure 2:**
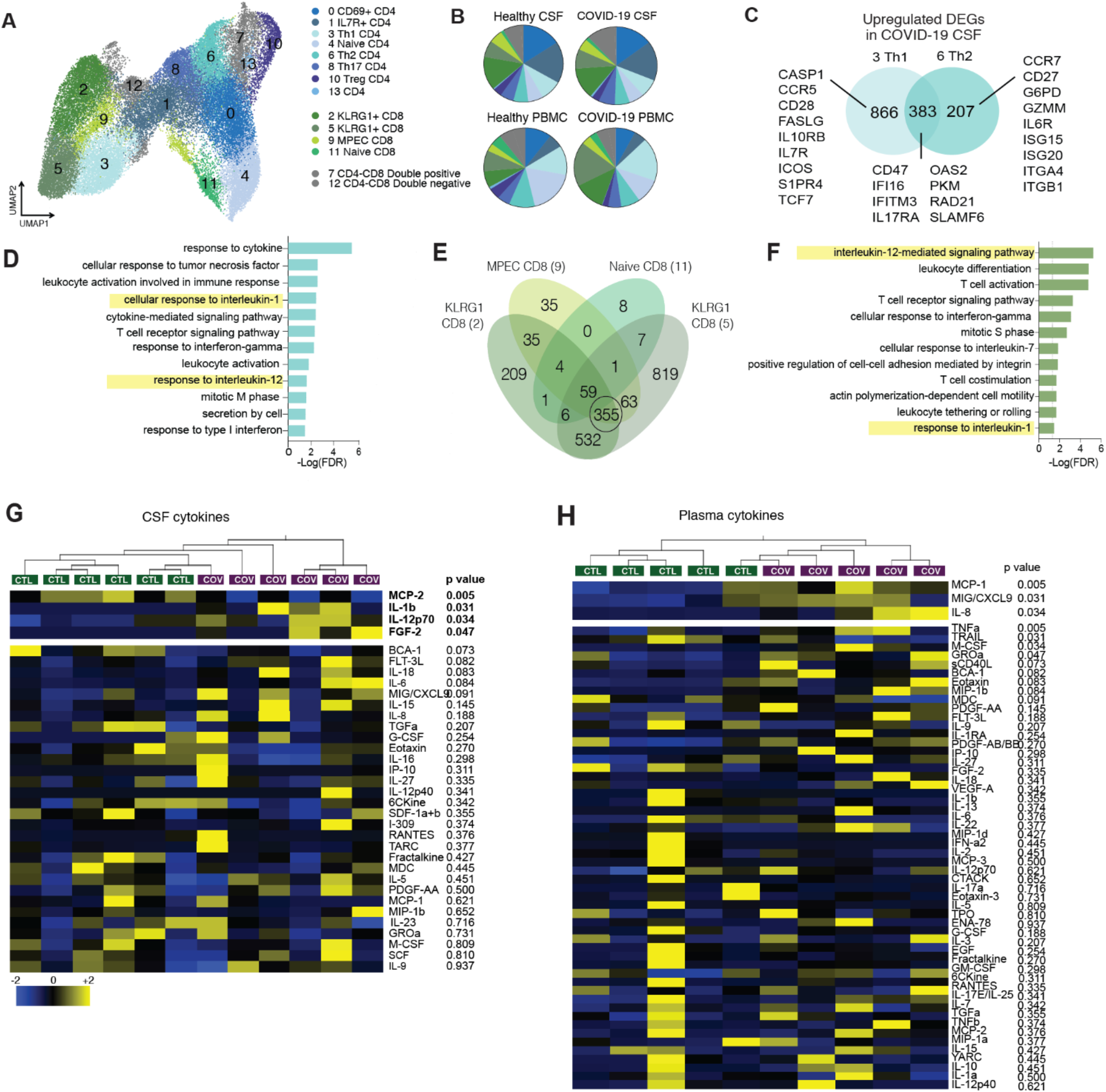
Transcriptional characterization of T cells in CSF and PBMC of COVID-19 patients. (**A**) Reclustered UMAP projection of combined CSF and peripheral blood T cells, demonstrating CD4 and CD8 T cell subsets. (Two KLRG1+ clusters are distinguished by GZMB and IFNG expression, see FigS3). (**B**) Pie charts depicting relative population frequency of different T cell subtypes found in CSF and PBMCs of control and COVID-19 patients. (**C**) Venn diagram depicting genes upregulated (adjusted p-value <0.05) in CSF of COVID-19 cases compared to PBMCs of COVID-19 cases in Th1 and Th2 CD 4 T cells. (**D**) Gene ontology analysis of genes that are upregulated in both Th1 and Th2 cells as depicted in Fig 2c. (**E**) Quad-Venn diagram of genes upregulated in CSF of COVID-19 patients compared to CSF of control patients in CD8 T cells. Genes shared by the three effector CD8 T cell subtypes are circled. (**F**) Gene ontology analysis of genes shared between the three effector CD8 T cell subtypes in Fig 2f. (**G** and **H**) Heat map of Luminex based cytokine profiling of CSF (G) and plasma (H) from COVID-19 cases and controls (n = 6 CSF, n= 5 plasma). For each cytokine, two-tailed *p* values were calculated using student’s t-test. Data for each row were mean-centered; each column shows data from one sample.

### Unique cytokine profiles exist in CSF of COVID-19 patients compared to serum

To validate the transcriptional enrichment in IL-12 and IL-1 signaling in the CSF of COVID-19 patients with neurologic symptoms, we measured inflammatory cytokine levels in the CSF and plasma using a Luminex cytokine panel (Figure 2g-h). Consistent with the single-cell RNA sequencing results, IL-1b and IL-12 were elevated in the CSF of COVID-19 cases compared to healthy controls, but were not elevated in the plasma of COVID-19 cases. Conversely, CCL2, CXCL9, and IL-8 were significantly increased in the periphery of COVID-19 cases compared to controls, but not in their CSF. Because IL-12 is thought to be produced by activated antigen presenting cells to orchestrate Th1 responses through T and NK cell activation, we examined the cellular source of IL-12 in COVID-19 cases. The innate immune cells with the highest *IL12A* expression were CSF macrophages, NK, and dendritic cells (Fig s1e). Taken together, these data support the single cell RNA sequencing analyses that identified IL-12 as differentially expressed in CSF but not blood innate immune cells of COVID-19 cases, and suggest a distinct effect of COVID-19 in the CNS on cytokines important for innate immunity and for the induction of cell-mediated immunity, including IL-1 and IL-12.

### CNS B cell responses to SARS-CoV-2 differ from those in the periphery

Having examined compartmentalized innate immune cells, T cells, and cytokine secretion, we next evaluated B cell populations in COVID-19 patients with neurologic symptoms. In these patients, we found a significant enrichment of B cells in the CSF when compared to CSF of controls (Figure 3a). Single cell RNA sequencing revealed several subtypes of B cells amongst PBMCs and in the CSF (Figure 3b, sFig 5a-c), including distinct plasma cell clusters. We therefore asked whether antibody-secreting B cells in the CSF exhibit a different anti-SARS-CoV-2 antibody profile than the those in the periphery. To do so, we utilized a recently developed SARS-CoV-2 epitope Luminex panel(*12*) to screen for anti-SARS-CoV-2 antibodies in the CSF and plasma of COVID-19 cases and controls. As expected, anti-SARS-CoV-2 antibodies were not detected in any controls. In contrast, all COVID-19 cases had anti-SARS-CoV-2 antibodies in both CSF and plasma. However, while all COVID-19 cases developed antibodies to SARS-CoV-2 spike and nucleocapsid in both the plasma and CSF, anti-RBD antibodies were rare in CSF but uniformly present in the plasma (Figure 3c). In addition, we found that in all COVID-19 cases, both the relative prevalence (rank score: 12 being most frequent, 1 being least frequent; Figure 3d) and levels of antibody (sFig 5d) diverged between the CSF and plasma, revealing a different anti-SARS-CoV-2 antibody profile between the CSF and plasma of the same patient.

**Figure 3:**
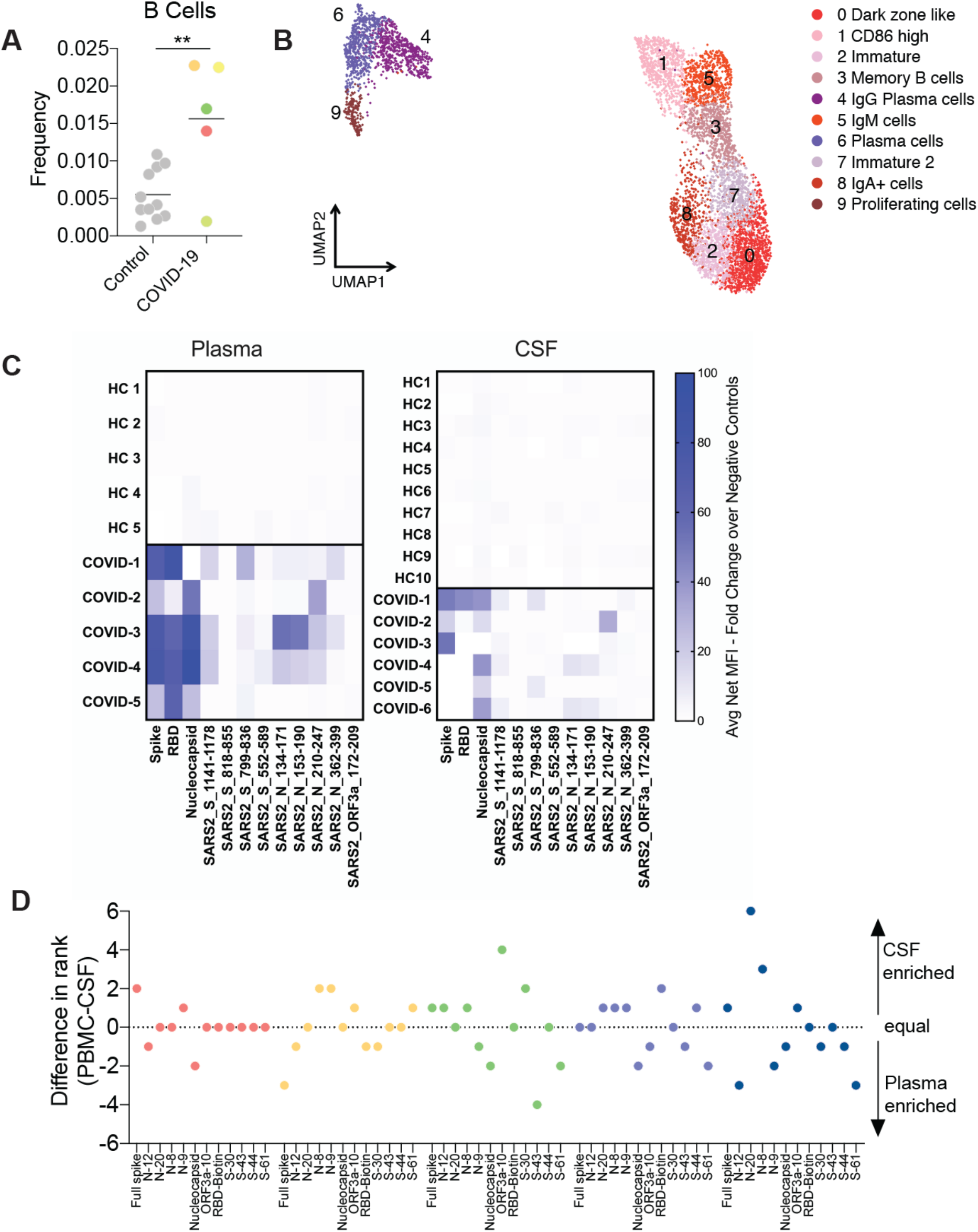
Localized central nervous system B cell responses in COVID-19 patients. (**A**) Frequency of B cells as a percentage of all CSF cells, in control and COVID-19 cases. (**B**) Re-clustered UMAP projection of B cells from CSF and blood (**C**) Heat map showing antibody binding in CSF (left) and plasma (right) to nine peptides from immunogenic regions of S, N and ORF3a, as well as whole S and N protein along with receptor binding domain of S protein. All data represented as fold change of fluorescent anti-IgG antibody signal over intra-assay negative controls. HC= healthy control. (**D**) Epitope frequency was ranked in each patient sample individually, and a difference in rank number for each cluster was graphed to determine CSF (positive) or Plasma (negative) enriched antibody epitopes. Two-tailed unpaired t-test **, p<0.01.

### A mouse model of CNS SARS-CoV-2 infection demonstrates compartmentalized CNS antibody secretion in response to CNS infection

Direct detection of SARS-CoV-2 in the CSF is extremely rare in reported cases of neurological complications of COVID-19(*13*), and SARS-CoV-2 RNA was similarly absent in our cohort. However, we detected intrathecal anti-viral antibodies in all cases. To determine whether the presence of CSF anti-SARS-CoV-2 antibodies in the absence of viral nucleic acid is consistent with viral neuroinvasion as is the case for other encephalitis-causing viruses including West Nile virus, Japanese encephalitis virus and measles virus(*14–16*), we used a recently developed mouse model that reliably dissociates pulmonary and neurological infection of SARS-CoV-2(*17*).

In our mouse model, adeno-associated virus is used to express the human ACE2 (hACE2) receptor in the lung, brain, or the lung and brain and allows us to target SARS-CoV-2 infection to specific tissues (sFig 6a-b). First, we used mice that express hACE2 in both the lung and the brain and administered SARS-CoV-2 intranasally (Figure 4a). This allowed us to establish SARS-CoV-2 infection in both the lung and the brain. In these mice, as expected, we detected increased titers of SARS-CoV-2 RNA in both the lung and brain tissue following inoculation. However, despite robust brain infection, we do not detect SARS-CoV-2 RNA in the CSF of these mice (Figure 4b). This suggests that direct detection of SARS-CoV-2 RNA in CSF at a single time point may be insensitive to parenchymal or short lived SARS-CoV-2 neuroinvasion.

**Figure 4:**
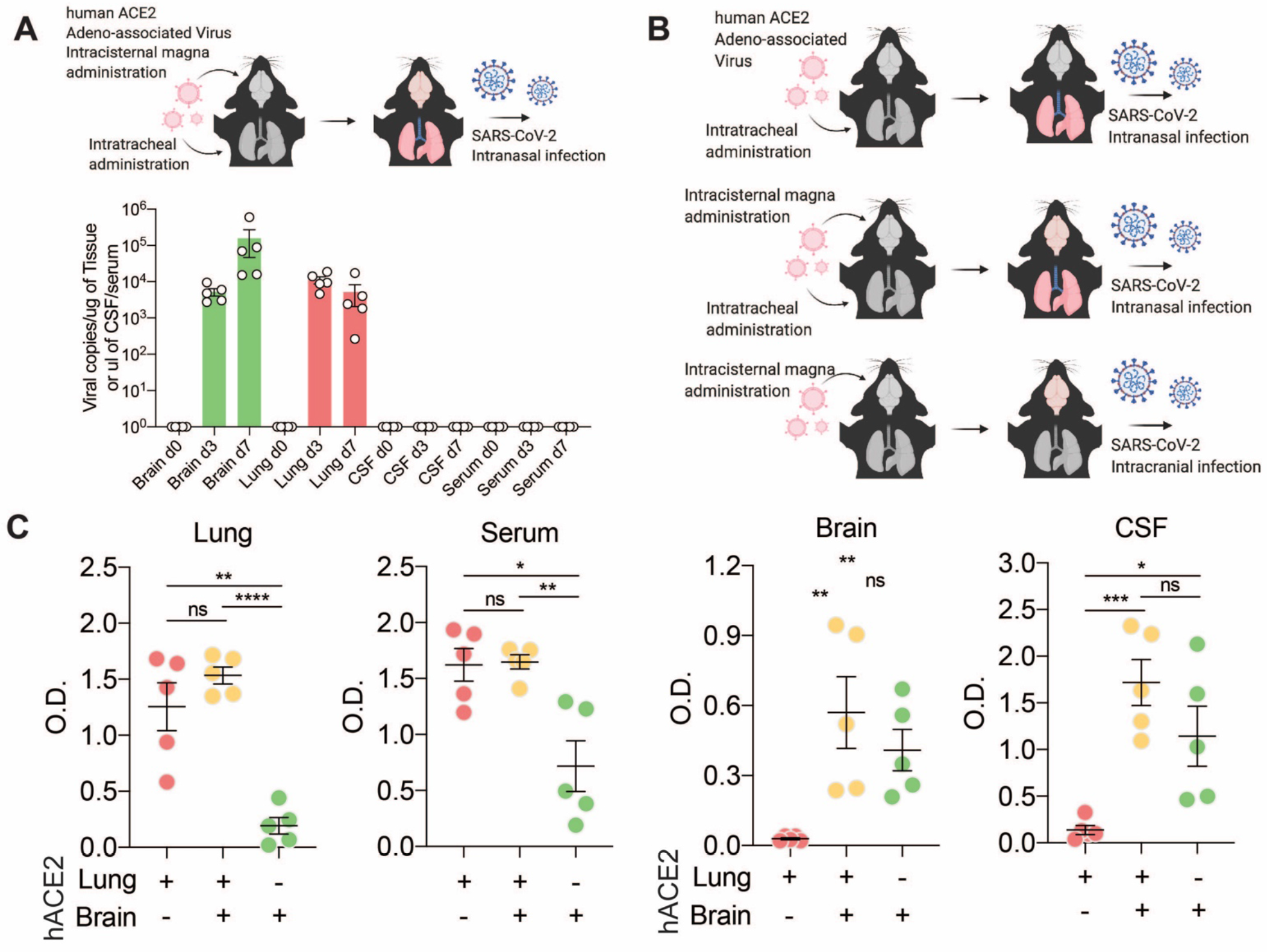
Cerebrospinal fluid antibodies reflect localized central nervous system infection. **(A)** Mice were transduced with AAV-hACE2 intrathecally and intratracheally for expression in the brain and lung and SARS-CoV-2 was introduced intranasally to establish brain and lung infection. Mice brains, lungs, CSF and serum collected at day 0 (before infection) and days 3 and 7 post infection and qPCR was performed to detect SARS-CoV-2 RNA. (**B**) Schematic of experimental procedure for (**C**). Mice were given localized AAV-hACE2 to overexpress human ACE2 in either the lung (top), brain and lung (middle) or brain only (bottom). After two weeks, mice were infected with SARS-CoV-2. (**C**) ELISA against SARS-CoV-2 spike protein was performed with lung homogenates, serum, brain homogenates and CSF. n = 5 for all three conditions. Two-tailed unpaired t-test (*, P<0.05; **, P<0.01; ***, P<0.005; ****, P<0.0001) and one-way ANOVA were performed (Lung, P<0.0001; Serum, P=0.002; Brain, P=0.0082; CSF, P=0.0016).

We next used the mouse model to evaluate whether the detection of intrathecal anti-SARS-CoV-2 antibodies in our COVID-19 patients was more likely triggered by a local antigen (i.e., as a consequence of SARS-CoV-2 neuroinvasion) or reflected passive transfer of antibody from the systemic circulation.(*13*) When SARS-CoV-2 was administered intranasally to mice expressing hACE2 only in the lung, we detected significantly elevated anti-spike SARS-CoV-2 IgG in the lung and serum of mice, while we saw little antibody signal in the brain or CSF of mice (Figure 4b, red). When SARS-CoV-2 was administered intranasally to mice expressing hACE2 in both the brain and lung, we detected increased anti-spike antibodies in all four compartments: lung, serum, brain, and CSF (Figure 4b, orange). Finally, when hACE2 was expressed in brain only and SARS-CoV-2 was administered intracranially (causing infection in the brain but not the lung), we detected increased anti-spike antibodies in the brain and CSF, but not in the lung or serum of the mice (Figure 4b, green). These data support that systemic infection alone may not be sufficient to account for the presence of humoral anti-viral immunity in the CNS. Indeed, in these mice, anti-spike antibodies in the CSF and brain were only observed in the setting of CNS infection, independent of whether there was an accompanying systemic SARS-CoV-2 infection.

#### Monoclonal antibodies from COVID-19 patient CSF are self-reactive

Although our COVID-19 cases did not have detectable SARS-CoV-2 RNA in their CSF, our CSF serology data suggested that CSF-expanded B cell populations might be specific for SARS-CoV-2 antigen(s). To address this question, we cloned individual monoclonal antibodies from the COVID-19 case with the largest number of clonally expanded B cell receptor (BCR) sequences in the CSF (n=5) and blood (n=4) (Case 1; Supplementary file 1). In this patient, the most prevalent BCR sequences comprised ∼25% and ∼10% of the total B cells in the blood and CSF, respectively (Figure 5a). Notably, the most prevalent BCR sequences detected in the CSF of case 1 did not overlap with the most prevalent peripheral BCR sequences (Figure 5b), supporting the hypothesis that a subset of CSF antibodies target antigen within the CNS. We found that one of five CSF-derived and two of four PBMC-derived monoclonal antibodies targeted SARS-CoV-2 spike protein (Figure 5c). None of the other monoclonal antibodies recognized other SARS-CoV-2 antigens on the Luminex panel. Given reports of new-onset humoral autoimmunity in COVID-19, we wondered whether some of the CSF-derived monoclonal antibodies might be autoreactive to neural tissue. Therefore, we tested all monoclonal antibodies for anti-neural immunoreactivity by anatomic mouse brain tissue staining. Monoclonal antibodies were used as a primary antibody to immunostain mouse brain tissue and labeled with an antihuman secondary antibody. An anti-influenza antibody targetting the hemagglutinin antigen (anti-HA) was used as a negative control (anti-INF). Similar to anti-HA, none of the PBMC-derived monoclonals recognized mouse brain tissue. In contrast, four of five CSF-derived monoclonal antibodies exhibited some degree of anti-neural immunoreactivity—including the anti-spike monoclonal (mAb C2) (Figures 5D and E). Notably, mAb C2 produced a neuropil-predominant immunostaining pattern on mouse brain tissue suggesting that antigen may be enriched in neuronal process or harbor an extracellular epitope. Immunoprecipitation mass spectrometry of mouse brain lysate using mAb C2 significantly enriched synaptopodin, a postsynaptic dendritic protein, and neural cadherin, a brain-enriched transmembrane protein that mediates presynaptic to postsynaptic adhesion. These data indicate that some CSF-expanded B cell populations in COVID-19 patients with neurologic symptoms are reactive to SARS-CoV-2 antigen, self-antigen, or both.

**Figure 5.**
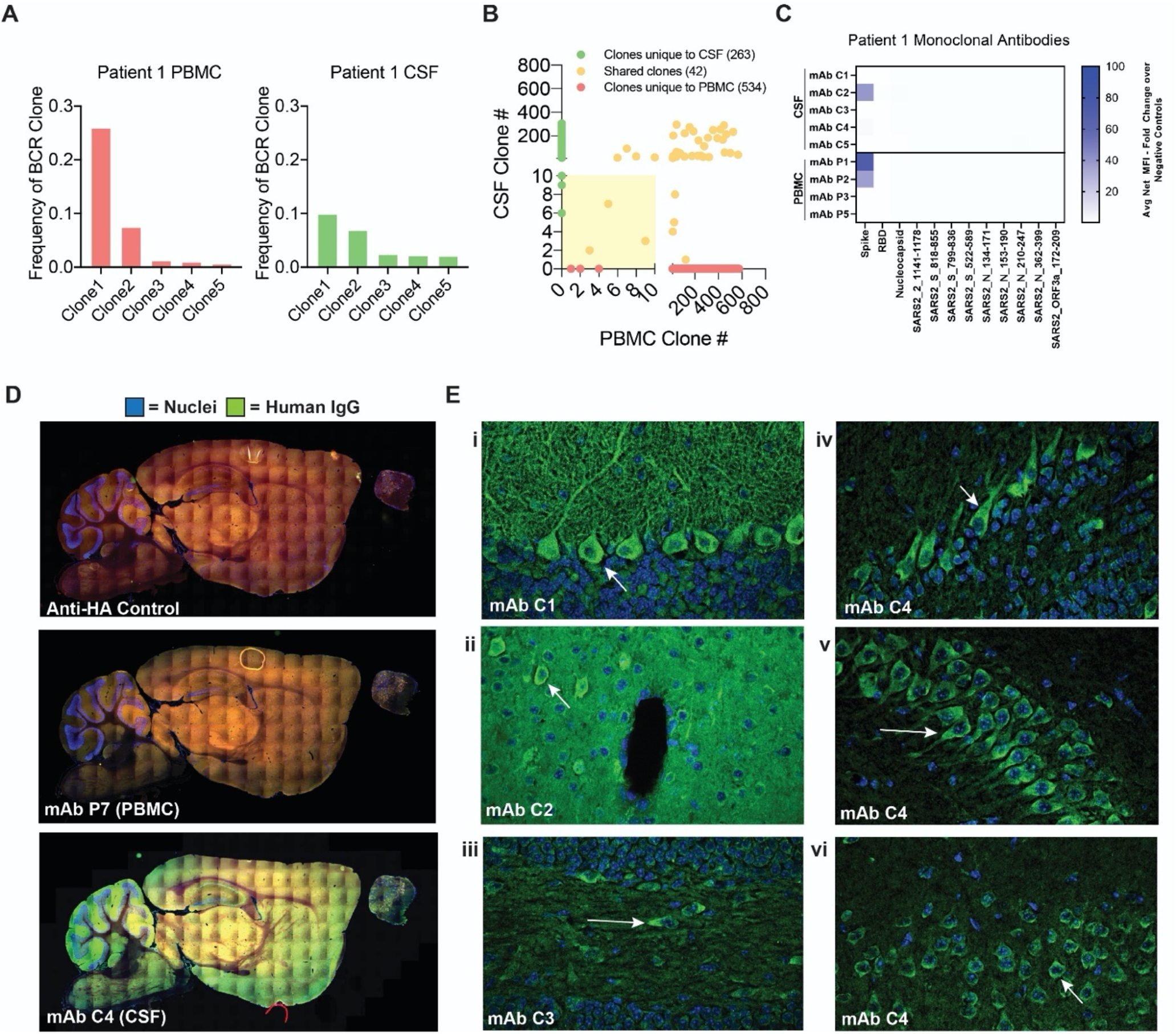
Antigenic Specificity of CSF- and PBMC-derived Monoclonal Antibodies. (**A**) Bar graph depicting frequency of top five most expanded clones in PBMC and CSF of patient 1. Bottom text shows uniquely distinct sequences in the top clones of this patient (shown in red). (**B**) Graph depicting overlap of clones found in CSF and PBMC of a patient. Green depicts clones only found in CSF, orange clones shared between the CSF and PBMC, and red, clones unique to PBMC. Yellow box depicts clones that would fall under top 10 most frequent clones in each compartment (lower clone rank # describes more frequent clone). (**C**) Heat map showing CSF-derived (mAbs C1 – C5) and PBMC-derived (mAbs P1 – 3 and P5) monoclonal antibody binding to nine peptides from immunogenic regions of S, N and ORF3a, as well as whole S and N protein along with receptor binding domain of S protein. Monoclonal antibody number corresponds to clone # from panels A and B; PBMC clone #4 (mAb P4) did not express well as a monoclonal antibody and was not used for subsequent studies. All data represented as fold change of fluorescent anti-IgG antibody signal over intra-assay negative controls. (**D**) Sagittal mouse brain sections were immunostained with mAbs 1 – 9 and a representative whole brain sagittal image is shown for PBMC-derived monoclonals (mAb 7) and CSF-derived monoclonals (mAb 4). An anti-hemagglutinin antibody (Anti-HA) in the same IgG1 backbone was used as a negati ve control. (**E**) Select regions of immunostaining from mAbs 1 – 4. Subpanels: i) mAb 1 immunostaining of cerebellar Purkinje cells (arrow) and the overlying molecular layer, ii) mAb C2 immunostaining of cortical neuropil and occasional staining of neuron-like soma (arrow), iii) mAb C3 immunostaining of large cells within the hilus of the hippocampus, iv) mAb C4 immunostains of mitral-like cells of the olfactory bulb (arrow), v) mAb C4 immunostaining of pyramidal neurons (arrow) in CA3 of the hippocampus vi) mAb C4 immunostaining of neuronal cell bodies in layer II of the cortex (arrow)

### Intrathecal humoral autoimmunity in COVID-19 patients with neurologic symptoms

The emergence of inflammatory and humoral autoimmune disorders of the nervous system during the para- or post-infectious period in COVID-19 is increasingly recognized and includes: acute disseminated encephalomyelitis (ADEM), autoimmune encephalitis associated with known autoantibodies, transverse myelitis, Guillain-Barré syndrome and one of its variants, Miller Fisher syndrome(*18–22*). Given this literature, and the autoreactivity of CSF-derived monocloncal antibodies from case 1, we hypothesized that our other COVID-19 cases might harbor intrathecal autoantibodies. To test this, we screened our cohort of COVID-19 cases for intrathecal anti-neural antibodies using a suite of complementary autoantigen detection platforms: anatomic mouse brain tissue immunostaining, immunoprecipitation mass spectrometry (IP-MS), and pan-human proteome phage display immunoprecipitation sequencing (PhIP-Seq)(*23–25*). In these screens we included one additional patient with post-COVID-19 seizures and cognitive impairment who had been recruited after the completion of the transcriptomic and cytokine analyses were completed (case 7, Supplementary Table 1, Supplementary File 1). Like the other six acute COVID-19 cases, this subject had detectable SARS-CoV-2 antibodies in his CSF (Supplementary Figure s7).

### Anatomic mouse brain immunostaining demonstrates the presence of intrathecal anti-neural autoantibodies in most COVID-19 cases

More COVID-19 CSF (5 of 7) were immunoreactive to mouse brain tissue at a 1:10 dilution than control CSF (2 of 6) (Figure 5A and Supplemental Figure s8). Control CSF staining was not specific to any anatomic region, weakly pan-nuclear, or primarily subpial (Supplemental Figure s9). None of the control CSF samples were immunoreactive beyond a 1:10 dilution, indicating the absence of high titer or high affinity anti-neural autoantibodies. In contrast, at a 1:10 dilution, COVID-19 CSF produced immunoreactive staining of specific anatomic regions including: cortical neurons (n = 4), the olfactory bulb (n = 3), thalamus (n = 3), CA3 field of the hippocampus (n = 3), cerebellum (n = 3), the brainstem (n = 4), and cerebral vasculature (n = 2) (Figure 5B & C, Supplemental Figure s8). Four and three COVID-19 CSF samples showed continuing immunoreactivity at 1:25 and 1:50 dilutions, respectively. The data indicate that an unexpectedly high proportion of CSF samples from COVID-19 patients with neurologic impairment harbor high titers of anti-neural autoantibodies of unknown pathogenic significance.

### IP-MS identifies intrathecal candidate autoantigens in a subset of COVID-19 cases

To screen for the neural protein targets of intrathecal autoantibodies, we immunoprecipitated whole mouse brain lysate using CSF and plasma, trypsinized precipitated proteins, and analyzed the resulting peptides by MS. IPs were performed in technical replicate, by different individuals, using different mice as input. First we searched resulting spectra against the SARS-CoV-2 proteome (Uniprot SARS-CoV-2 reference proteome, 06/2020). SARS-CoV-2 proteins were not detected. We then searched for human proteins. Consistent with circulating protein expression patterns(*26, 27*) and circulating prothrombotic autoantibodies(*28*) in COVID-19, complement, coagulation, and platelet degranulation pathway proteins of human origin were significantly overrepresented in the IgG-bound protein fraction of plasma from our cases (fig. S10).

To specifically identify candidate anti-neural autoantibodies in the CSF, we searched for mouse proteins that were observed in both technical replicates and significantly enriched by spectral counting and/or 1.5x enriched by MS1 peak area in COVID-19 CSF IPs relative to controls. By IP-MS, between 5 and 56 (median = 20) proteins were enriched by the CSF, but neither the plasma of the same case nor control CSF, indicating they were unique to the CSF compartment of COVID-19 patients. By gene ontology pathway analysis, the top three cellular components that IP-MS CSF-specific proteins mapped to were the myelin sheath (e.g. CNP and NSF), post-synaptic density proteins (e.g. PSD-95 and SYN2), and the postsynaptic cytoskeleton (e.g. INA and MYO5A) (Bonferroni-corrected p = 1.3 × 10^−17^, 3.3×10^−8^, and 1.6×10^−6^ respectively).

### PhIP-Seq identifies intrathecal candidate autoantigens in a subset of COVID-19 cases

COVID-19 CSF was also screened for autoantibodies using a previously described PhIP-Seq platform (T7 bacteriophage display) displaying ∼730,000 overlapping 49 amino acid peptides spanning all human proteins, including all known and predicted isoforms(*24, 25*). To identify peptides that were significantly enriched by COVID-19 CSF compared to controls, we first established an empirical enrichment threshold using a validated commercial antibody targeting the protein GFAP (Supplemental File 2). For COVID-19 CSF samples, peptides with suprathreshold enrichment in both technical replicates were considered candidate autoantigens. COVID-19 CSF enriched between 2 and 40 CSF-specific proteins (median = 18, Supplementary File 2). By gene ontology, synaptic proteins were also enriched in CSF-specific PhIP-Seq candidate autoantigens (e.g., NRG3, SYNJ2, and DPYSL2) (Bonferroni-corrected p = 1.6x×10^−1^). COVID-19 CSF that was immunoreactive to mouse brain tissue at a 1:50 dilution was associated with greater enrichment of candidate autoantigens by PhIP-Seq (19 – 40, median = 37) than CSF that immunostained at a lower dilution or was not immunoreactive (1 – 16, median = 6) suggesting a correlation between immunostaining status and the burden of CSF autoantibodies. In some instances, the same candidate autoantigen was detected in COVID-19 cases by IP-MS and PhIP-seq: UHRF1BP1 (Case 2), NUAK1 (Case 3), and DBN1 (Case 7).

### HEK293 overexpression cell-based assay validates two candidate autoantigens in the CSF of COVID-19 cases

To validate anti-neural autoantibodies identified in CSF, we selected two COVID-19 candidate autoantigens that were enriched by CSF but not plasma on PhIP-Seq: intraflagellar transport protein 88 homolog (IFT88) and syntaphilin (SNPH). We validated them by HEK293 overexpression cell-based assay (CBA) (Figure 5D). HEK293 cells were transfected with a plasmid encoding the candidate autoantigen, fixed, and immunostained with patient CSF. By CBA, both anti-IFT88 and anti-SNPH validated as *bona fide* autoantibodies in the CSF of COVID-19 Cases 1 and 3, respectively, but not control CSF (Figure 6E and Supplementary Figure s11). Taken together, these autoantibody data indicate that: a high proportion of COVID-19 CSF is immunoreactive to neural tissue, immunoreactive CSF enrich more CSF-specific candidate autoantigens than non-immunoreactive CSF, and some candidate autoantigens represent *bona fide* autoantibodies as verified by CBA.

**Figure 6:**
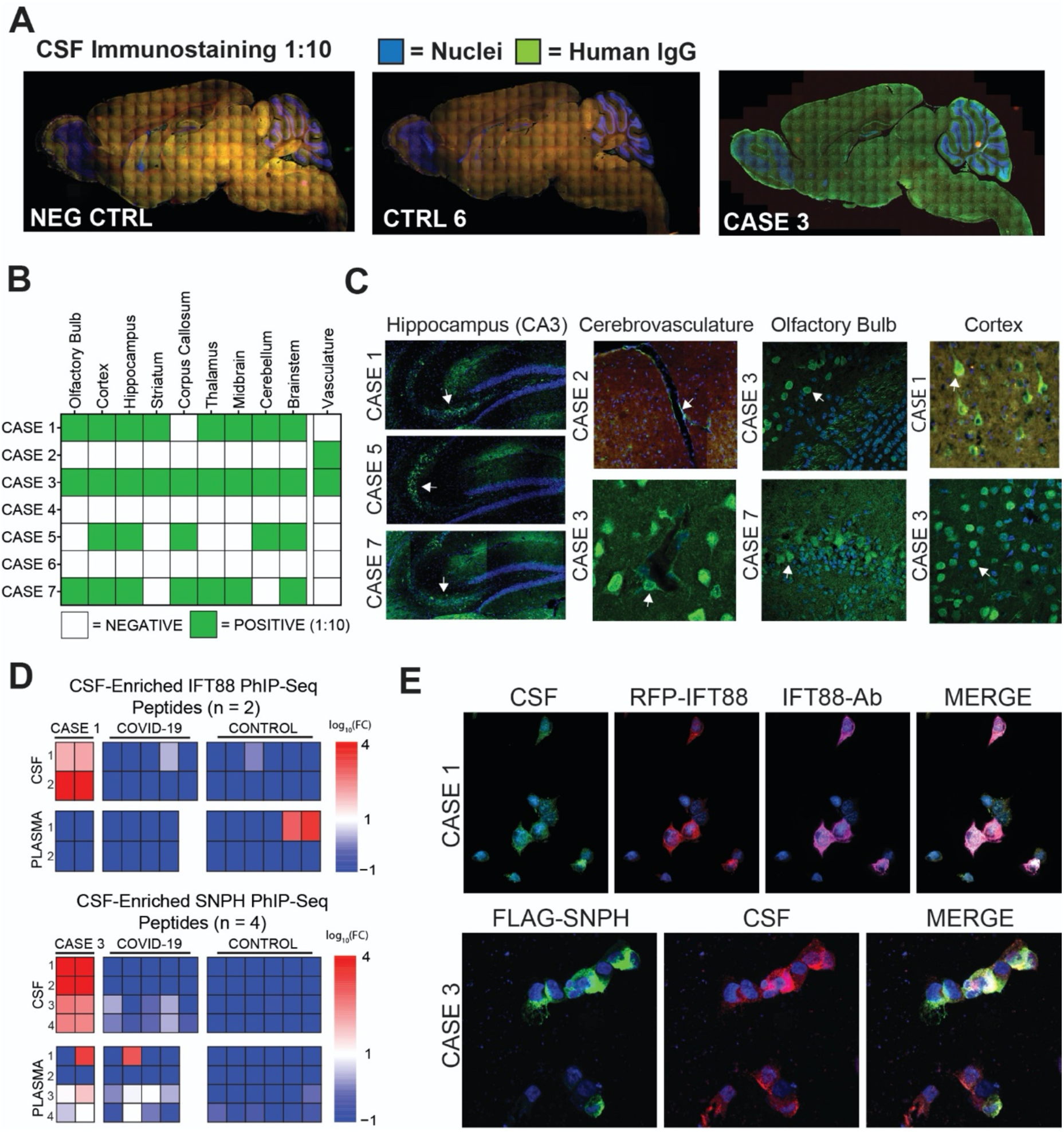
Autoantibodies in the CSF of COVID-19 patients. **(A)** Sagittal mouse brain sections were immunostained with CSF at a 1:10 dilution (green) and the nuclear stain DAPI (blue). Secondary only negative control (left) and 4/6 control CSF (example CTRL 6, center) were not immunoreactive. In contrast, 5/7 COVID-19 CSF samples were immunoreactive (example CASE 3, right) **(B)** Binary matrix indicating anatomic immunoreactivity of COVID-19 CSF at a 1:10 dilution. **(C)** Select examples of COVID-19 CSF anatomic immunostaining: CA3 of the hippocampus (n = 3, arrows, left column), cerebrovasculature (top panel second column arrow shows endothelial staining; bottom panel arrow shows a perivascular cell), olfactory bulb (n = 3, two shown, third column, top panel neuron-like cells; bottom panel mitral cells), cortical neuron-like cells (n = 4, two cases shown, fourth column). **(D)** Heatmaps of sequence-sharing peptides mapping to IFT88 (Case 1, top) and SNPH (Case 3, bottom) that were enriched by CSF shown with their corresponding enrichment by plasma. Row = values for individual peptides. Left two columns = individuals for retchnical replicates for case 1 (upper) and case 3 (lower). COVID-19 and CONTROL columns = the mean of the log_10_(fold change enrichement) for individual cases. For both case 1 and case 3, candidate IFT88 and SNPH peptides, respectively, were enriched by CSF but not plasma. **(E)** HEK 293 overexpression cell-based assays demonstrating that Case 1 CSF (top, green) is immunoreactive to overexpressed IFT88 (top, red) and Case 3 CSF (bottom red) is immunoreactive to overexpressed SNPH (bottom green).

## Discussion

In this exploratory study, we performed a comprehensive set of immunologic investigations to assess for CNS-specific immune responses in a series of COVID-19 patients with neurologic symptoms(*29, 30*). While systemic multi-organ dysfunction is almost certainly a driver of neurologic complications in a proportion of COVID-19 patients, here we identified both innate and adaptive anti-viral immune responses, as well as humoral autoimmunity that appears to be unique to the CNS, and may therefore contribute to COVID-19 neuropathology.

CSF, while not identical to brain, is produced by the choroid plexus and bathes the CNS. It is the only CNS tissue surrogate readily sampled in living humans. Analysis of CSF immune cells has shed light on immune mechanisms of neuronal injury during other infections, including HIV, neurosyphilis, and neuroborreliosis(*31–33*). By assessing CSF and blood in patients with acute COVID-19 and neurological symptoms, we find evidence for a compartmentalized CNS immune response to SARS-CoV-2. Through transcriptional and cytokine analyses, we find an increase in CSF but not plasma IL-12 and IL-1b, factors that are central for coordinating innate and adaptive immune responses to invading pathogens. Notably, mouse CoV neuroinvasion also leads to IL-12 production by astrocytes and microglia(*34*).

Our data identified increased and divergent humoral responses within the CNS. This humoral response included indirect evidence for neuroinvasion of SARS-CoV-2 through the presence of anti-viral antibodies in the CNS during acute SARS-CoV-2 infection. Indeed, using an animal model, we show that SARS-CoV-2 infection in the CNS is necessary for stimulating the production of intrathecal SARS-CoV-2 antibodies as isolated systemic viral infection is associated with little to no antibody in CSF. These data therefore suggest that the presence of anti-SARS-CoV-2 antibody in the CSF of COVID-19 patients may similarly reflect viral infection of the CNS. Further supporting this contention, we generated a monoclonal antibody from the CSF of a COVID-19 patient (case 1) with marked B cell clonal expansion in the CSF that was specific for the SARS-CoV-2 spike protein. This B cell clone was not detected in the peripheral blood of the same patient. Notably, the other CSF-derived monoclonal antibodies were variably immunoreactive to mouse brain tissue which motivated and expanded search for autoantibodies in the CSF of our other COVID-19 cases.

Indeed, our extended studies of the CNS humoral response suggest that a subset of COVID-19 patients with neurologic symptoms have an elevated burden of autoreactive antibodies in their CSF. By anatomic immunostaining, an unexpectedly high proportion of our COVID-19 cases harbored intrathecal autoantibodies, including anti-neural autoantibodies. Hippocampal immunostaining from three cases was primarily restricted to the CA3 region similar to a recent report of SARS-CoV-2 associated encephalitis(*35*). Other immunoreactive anatomic regions included the olfactory bulb in three cases and cerebrovasculature in two— anatomic regions with *prima facie* relevance to common neurologic sequelae of COVID-19 (i.e., anosmia and stroke). Subsequent unbiased protein and peptide screens of CSF identified a diversity of candidate autoantibodies, and two of these autoantigens, SNPH and IFT88, were subsequently validated by CBA. Although SNPH and IFT88 are previously undescribed autoantigens of indeterminate clinical significance, their biologic functions suggest they could be implicated in autoantibody-mediated neurologic dysfunction. SNPH is a brain-enriched transmembrane protein that inhibits SNARE complex formation and subsequent neurotransmitter release(*36*). IFT88 is a ciliary protein whose mutation causes a ciliopathy in humans and anosmia in mice(*37*).

However, the mere presence of an intrathecal autoantibody does not mean that a patient has an autoimmune encephalitis. Indeed, the patients in our exploratory cohort lacked evidence for active inflammation on neuroimaging and/or did not have elevated conventional CSF markers of neuroinflammation (i.e., white blood cell count, IgG index, and CSF-restricted oligoclonal bands) typically, but not always, found in patients with autoimmune encephalitis. Moreover, our study is complicated by the diverse range of neurological symptoms in our study participants. Finally, a comparison group with a different systemic viral infection would help determine which aspects of the findings are specific to COVID-19. Nonetheless, our data suggest that deeper CNS phenotyping of COVID-19 patients with acute and/or chronic neurologic sequelae should be pursued. Notably, COVID-19 patients with neurological symptoms appear to have immune responses to multiple autoantigens, implying that the increased compartmental humoral immune response may reflect a broader immune activation syndrome. This is particularly true given that humoral autoimmunity has been observed to target other organ systems in COVID-19, and may also contribute to neuropathology during COVID-19(*38–40*),(*41*).

Taken together, our exploratory data suggest that even in COVID-19 patients with neurologic symptoms who lack overt evidence for neuroinflammation on MRI or conventional CSF testing, there is a compartmentalized immune response involving the innate and adaptive arms of the immune system. Future research, involving careful clinical phenotyping and timely investigations of the CSF, will help place these findings into a broader clinical context and inform whether anti-viral and/or immunomodulatory therapies might be indicated for carefully selected neurologically impaired patients with COVID-19.

## Materials and Methods

All procedures were performed in a BSL-3/ABSL-3 facility (for SARS-CoV-2-related work) with approval from the Yale Environmental Health and Safety committee (#20-19 and #18-16).

### Study participants

COVID-19 patients admitted to Yale New Haven Hospital in March-May 2020 were recruited to the IRB approved Yale IMPACT study (Implementing Medical and Public Health Action Against Coronavirus CT). COVID-19 patients who were undergoing clinical lumbar puncture for evaluation of neurological symptoms were included. Negative controls were recruited from the surrounding community via IMPACT and the IRB approved HARC study at Yale. Healthy control participants recruited after March 2020 were confirmed to be negative for SARS-CoV-2 infection by nasopharyngeal swab PCR. All participants or their designated surrogate consented to donate 25cc of CSF for research studies. Blood was collected within one hour of lumbar puncture. For autoantibody characterization assays, additional pre-pandemic healthy volunteers (CTRLS 7 - 12, supplemental table S2) were recruited from the general population via word of mouth, newspaper and internet advertisements, and poster flyers. Based on a questionnaire, they were excluded if they had an Axis I psychiatric diagnosis, a first-degree relative with a known or suspected Axis I psychiatric disorder, or any neurologic disorder. CSF samples were collected at The Zucker Hillside Hospital, Glen Oaks, NY.

### SARS-CoV-2 RT-qPCR

RNA was extracted from nasopharyngeal swabs, CSF, and plasma using the MagMax Viral/Pathogen Nucleic Acid Isolation kit. A modified CDC RT-qPCR assay was used to detect SARS-CoV-2 with the N1, N2, and human RNase P (RP) primer-probe sets and the NEB Luna Universal Probe One-Step RT-qPCR kit on the Bio-Rad CFX96 Touch Real-Time PCR Detection System(*8*).

### PMBC and CSF cell preparation

Peripheral blood mononuclear cells (PBMCs) were isolated from heparinized whole blood after 1:1 PBS dilution. The blood was layered over a Histopaque (Sigma-Aldrich, #10771-500ML) gradient in a SepMate tube (Stemcell Technologies; #85460) and isolated according to manufacturer’s instructions. The PBMCs were then aliquoted and stored at −80 for subsequent analysis. CSF was centrifuged at 400G for 8 minutes, with cell-free supernatant removed for cytokine and antibody assays, and cell pellet processed for single cell RNA sequencing, as below.

### Single cell RNA sequencing

Approximately 8,000 single cells from CSF and from PBMC from each participants were loaded into each channel of a Chromium single-cell 5′ Chip (V3 technology). 5’ 10X libraries were sequenced on Illumina Novaseq at approximately 50,000 reads per cell. Raw sequencing reads were aligned to the human GRCh38 genome and gene counts were quantified as UMIs using Cell Ranger count v3.0 (10x Genomics). We removed cells with >10% mitochondrial RNA content, and included cells with > 500 and <2000 genes expressed per cell. Dimensionality reduction, clustering, and visualization was performed using Seurat. Interferon regulated genes were identified using Interferome(*42*). Differential expression analysis was performed in Seurat v3(*43*) using the two-tailed Wilcoxon test, comparing cells from COVID-19 patients vs healthy controls. Significant differentially expressed genes (DEGs) were defined as: adjusted *p* < 0.05 and |log fold change| ≥ 0.1. Gene ontology and pathway enrichment analysis was performed using DAVID(*44*). DEGs in PBMC and CSF samples were compared using the UpSetR package.

To identify potential intercellular interactions between different cell types in the scRNA-seq data, we utilized CellPhoneDB v2(*10*). Normalized count matrices and associated cell type labels were provided to CellPhoneDB and analyzed under both the statistical mode and the thresholding mode. Of note, since the statistical mode of CellPhoneDB seeks to assess the specificity of a given interaction, a lack of statistical significance does not necessarily mean a given interaction is not present. Therefore, when comparing the number of potential intercellular interactions in COVID-19 patients vs healthy controls, the simpler threshold-based analysis mode was used. In contrast, for pinpointing the top candidate cell-cell interactions in each dataset, the statistical analysis mode was used, with a significance threshold of *p* < 0.05.

To further explore the downstream consequences of these intercellular interactions, we performed NicheNet analysis(*11*). The cDC1, NK cell, monocyte, macrophage, and myeloid cell clusters were defined as the potential “sender” cell populations. Different lymphocyte populations were defined as potential “receiver” cells: all CD4 T cells merged across clusters, merged CD8 T cells, or merged B cells. Potential ligands and receptors were defined as being expressed in ≥ 20% of cells in a particular cell type. Differentially expressed genes (i.e. candidate target genes) were defined similarly as above, with adjusted *p* < 0.05 and |log fold change| ≥ 0.1.

### T and B cell clustering

We initially combined all 76,473 CSF and blood cells and generated clusters using Seurat. For each cluster we assigned a cell-type label using statistical enrichment for sets of marker genes, and manual evaluation of gene expression for small sets of known marker genes. We then created a separate Seurat object consisting only of T cells clusters from the original analysis, and a separate Seurat object consisting only of plasma and B cells. We then re-clustered these T and B cells and annotated sub-clusters using previously annotated marker genes.

### BCR analysis

Single cell V(D)J sequences were generated using CellRanger vdj function. Assignments of V(D)J sequences were performed using IgBLAST v.1.6.1 with the September 12, 2018 version of the IMGT gene database (as described previously)(*45, 46*). Non-functional V(D)J sequences were removed. Cells with multiple IGH V(D)J sequences were assigned to the most abundant V(D)J sequence by unique molecular identifier count (and based on numbers of sequenced reads in instances with ties). B cell clones in the CSF and circulation were identified using an approach described previously using hierarchical clustering implemented using the DefineClones.py function of Change-O v0.3.4 and a junctional sequence hamming dissimilarity threshold of 0.17(*46*). To account for the presence of light chains, heavy chain-based clones were corrected for using an approach described previously(*47*).

### Cytokine assays

Soluble chemokines and cytokines were assessed in CSF supernatant and paired plasma using the HD71 Human Cytokine Array/Chemokine Array (Eve Technologies, Calgary, AB). Statistical analysis was carried out using Qlucore Omics Software, version 3.6 (Lund, Sweden). Cytokines that were absent from CSF or plasma under both COVID and controls conditions were excluded from the respective analyses. Heatmaps were plotted using Z-scores, with the color scale set to range from −2 to +2. Hierarchical clustering was applied to samples.

### SARS-CoV-2 Serological Assay

Highly immunogenic linear regions of the SARS-CoV-2 proteome were isolated by ReScan and conjugated to Luminex beads as previously described(*48*). Briefly, high concentration T7 phage stocks displaying immunodominant epitopes of the S, N and ORF3a proteins were propagated and grown to high (>10^11^ PFU/mL) titer then were each conjugated to unique bead IDs according to manufacturer’s Antibody Coupling Kit instructions (Luminex). Whole N protein (RayBiotech) beads were conjugated similarly using manufacturer instructions with 5µg of protein per 1 million beads. For other whole protein Luminex-based beads, MagPlex-Avidin Microspheres (Luminex) were coated with either the S protein RBD (residues 328-533) or the trimeric S protein ectodomain (residues 1-1213).

All beads were blocked overnight before use and pooled on day of use. 2000-2500 beads per ID were pooled per incubation with patient plasma at a final dilution of 1:500 or patient CSF at a final dilution of 1:20 for 1 hour, washed, then stained with an anti-IgG pre-conjugated to phycoerythrin (Thermo Scientific, #12-4998-82) for 30 minutes at 1:2000. Primary incubations were done in PBST supplemented with 2% nonfat milk and secondary incubations were done in PBST. Beads were processed in 96 well format and analyzed on a Luminex LX 200 cytometer. Median Fluorescence Intensity from each set of beads within each bead ID were retrieved directly from the LX200 and log transformed after normalizing to the mean signal across two intra-assay negative controls (glial fibrillary acidic protein (GFAP) and tubulin phage peptide conjugated beads).

### Mice for SARS-CoV-2 model

Six to twelve-week-old mixed sex C57Bl/6 (B6J) purchased from Jackson laboratories were subsequently bred and housed at Yale University. All procedures used in this study (sex-matched, age-matched) complied with federal guidelines and the institutional policies of the Yale School of Medicine Animal Care and Use Committee.

### AAV infection (Intratracheal and Intracisternal magna injection)

Adeno-associated virus 9 encoding hACE2 were purchased from Vector biolabs (AAV-CMV-hACE2). *Intratracheal injection.* Animals were anaesthetized using a mixture of ketamine (50 mg kg^−1^) and xylazine (5 mg kg^−1^), injected intraperitoneally. The rostral neck was shaved and disinfected. A 5mm incision was made and the salivary glands were retracted, and trachea was visualized. Using a 500μL insulin syringe a 50μL bolus injection of 10^11^GC of AAV-CMV-hACE2 was injected into the trachea. The incision was closed with VetBond skin glue.

Following intramuscular administration of analgesic (Meloxicam and buprenorphine, 1 mg kg^−1^), animals were placed in a heated cage until full recovery. *Intracisternal magna injection.* Mice were anesthetized using ketamine and xylazine, and the dorsal neck was shaved and sterilized. A 2 cm incision was made at the base of the skull, and the dorsal neck muscles were separated using forceps. After visualization of the cisterna magna, a Hamilton syringe with a 15 degree 33 gauge needle was used to puncture the dura. 3μL of AAV_9_ (3.10^12^ viral particles/mouse) or mRNA (4-5 μg) was administered per mouse at a rate of 1μL min^−1^. Upon completion of the injection, needle was left in to prevent backflow for an additional 3 minutes. The skin was stapled, disinfected and same post-operative procedures as intratracheal injections were performed.

### Generation of SARS-CoV-2 virus

To generate SARS-CoV-2 viral stocks, Huh7.5 cells were inoculated with SARS-CoV-2 isolate USA-WA1/2020 (BEI Resources #NR-52281) to generate a P1 stock. To generate a working stock, VeroE6 cells were infected at a MOI 0.01 for four days. Supernatant was clarified by centrifugation (450g x 5min) and filtered through a 0.45 micron filter. To concentrate virus, one volume of cold (4 °C) 4x PEG-it Virus Precipitation Solution (40% (w/v) PEG-8000 and 1.2M NaCl) was added to three volumes of virus-containing supernatant. The solution was mixed by inverting the tubes several times and then incubated at 4 °C overnight. The precipitated virus was harvested by centrifugation at 1,500 x g for 60 minutes at 4 °C. The pelleted virus was then resuspended in PBS then aliquoted for storage at −80°C. Virus titer was determined by plaque assay using Vero E6 cells. *SARS-CoV-2 infection (intranasal):* Mice were anesthetized using 30% v/v Isoflurane diluted in propylene glycol. Using a pipette, 50μL of SARS-CoV-2 (3×10^7^ PFU/ml) was delivered intranasally. *SARS-CoV-2 infection (intraventricular):* Animals were anaesthetized using a mixture of ketamine (50 mg kg^−1^) and xylazine (5 mg kg^−1^), injected intraperitoneally. After sterilization of the scalp with alcohol and betadine, a midline scalp incision was made to expose the coronal and sagittal sutures, and a burr holes were drilled 1 mm lateral to the sagittal suture and 0.5 mm posterior to the bregma. A 10 μl Hamilton syringe loaded with virus, and was inserted into the burr hole at a depth of 2 mm from the surface of the brain and left to equilibrate for 1 min before infusion. Once the infusion was finished, the syringe was left in place for another minute before removal of the syringe. Bone wax was used to fill the burr hole and skin was stapled and cleaned. Following intramuscular administration of analgesic (Meloxicam and buprenorphine, 1 mg kg^−1^), animals were placed in a heated cage until full recovery. For high condition, 5μL of SARS-CoV-2 (3×10^7^ PFU/ml) and for low condition 5μL of SARS-CoV-2 (3×10^6^ PFU/ml) was used.

### Enzyme-linked immunosorbent assay

ELISAs were performed as previously reported(*49*). In short, Triton X-100 and RNase A were added to serum samples at final concentrations of 0.5% and 0.5mg/ml respectively and incubated at room temperature (RT) for 3 hours before use to reduce risk from any potential virus in serum. 96-well MaxiSorp plates (Thermo Scientific #442404) were coated with 50 μl/well of recombinant SARS Cov-2 S1 protein (ACROBiosystems #S1N-C52H3-100ug) at a concentration of 2 μg/ml in PBS and were incubated overnight at 4 °C. The coating buffer was removed, and plates were incubated for 1h at RT with 200μl of blocking solution (PBS with 0.1% Tween-20, 3% milk powder). Serum was diluted 1:50 in dilution solution (PBS with 0.1% Tween-20, 1% milk powder) and 100μl of diluted serum was added for two hours at RT. Plates were washed three times with PBS-T (PBS with 0.1% Tween-20) and 50μl of mouse IgG-specific secondary antibody (BioLegend #405306, 1:10,000) diluted in dilution solution added to each well. After 1h of incubation at RT, plates were washed three times with PBS-T. Samples were developed with 100μl of TMB Substrate Reagent Set (BD Biosciences #555214) and the reaction was stopped after 15 min by the addition of 2 N sulfuric acid.

### Statistical methods

All statistical analyses were performed using commercially available software (Prism or Excel). All values are expressed as the mean ± SEM. Differences in means between two groups were analysed using unpaired two-sided *t*-tests, unless otherwise noted. For scRNA-seq analyses, we corrected for multiple comparisons and report adjusted P values using Benjamini–Hochberg correction. For pathway analyses, Fisher’s exact test was used with Bonferroni correction for multiple testing.

### Anatomic Mouse Brain Tissue Staining

Postnatal day 40 – 60 mice (F1 generation of FVB x C57blk6 cross) were transcardially perfused with 4% paraformaldehyde (PFA) and brains post-fixed in PFA overnight. After sucrose equilibration, brains were blocked in OCT and sectioned at 12µm on a standard cryostat. Sections were permeablized and blocked in PBS containing 10% lamb serum and 0.1% triton x-100. Sections were then incubated with CSF at 1:10, 1:25, and 1:50 overnight at 4C. Sections were rinsed at least 5x with PBS and counterstained with anti-human IgG (Alexafluor 488). Nuclei were stained with DAPI at 1:2000 and stained sections were coversliped with prolong gold.

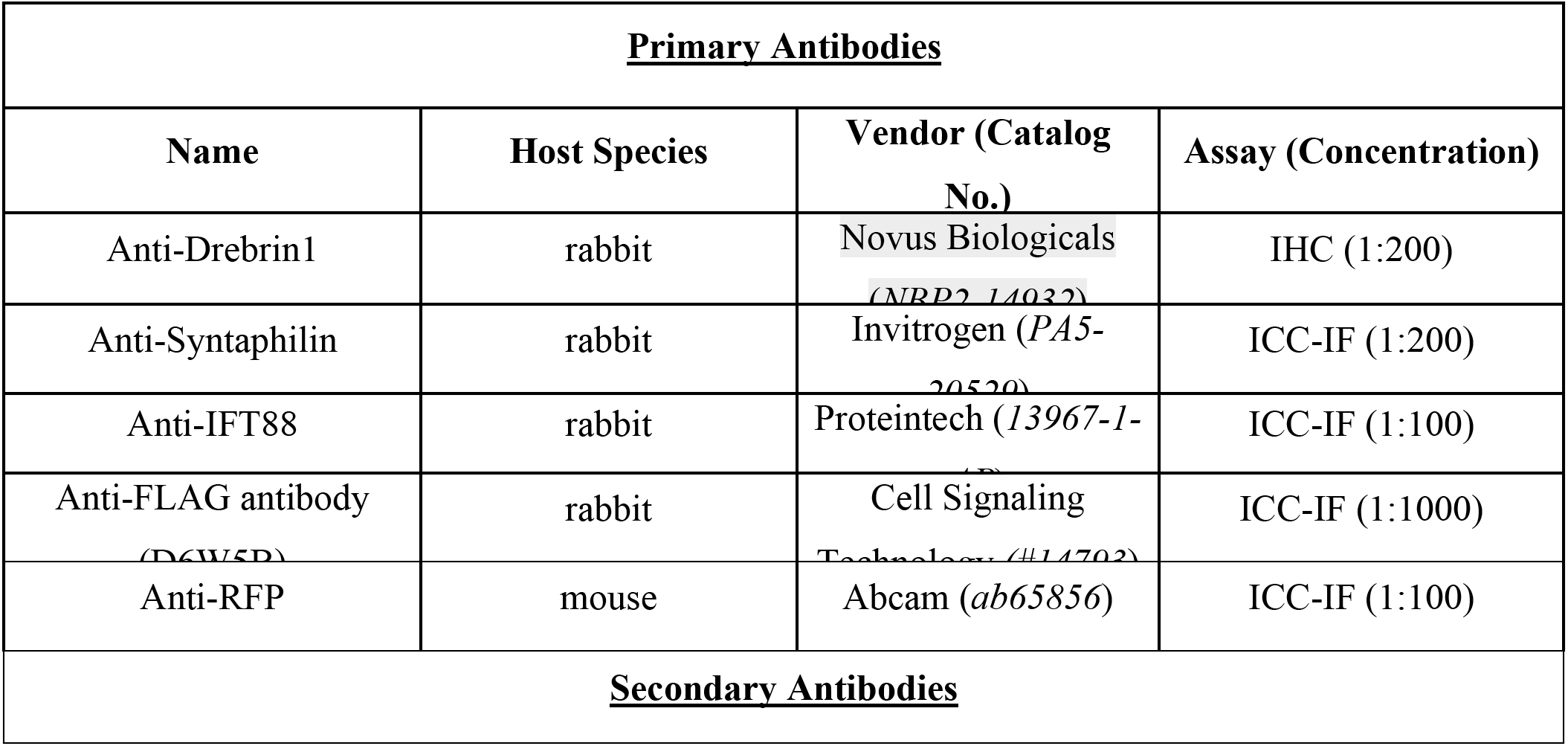
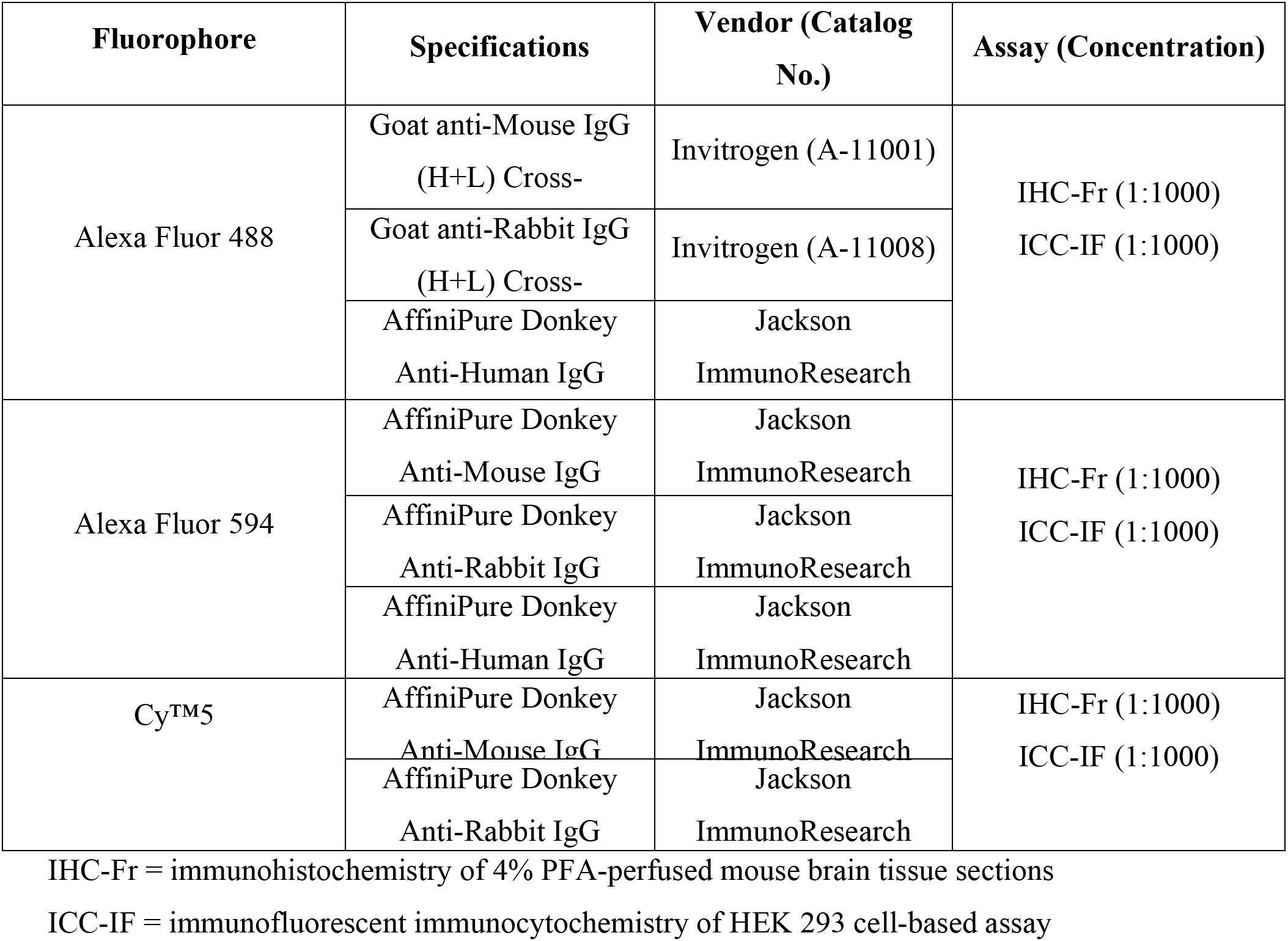

### HEK 293 Overexpression Assays

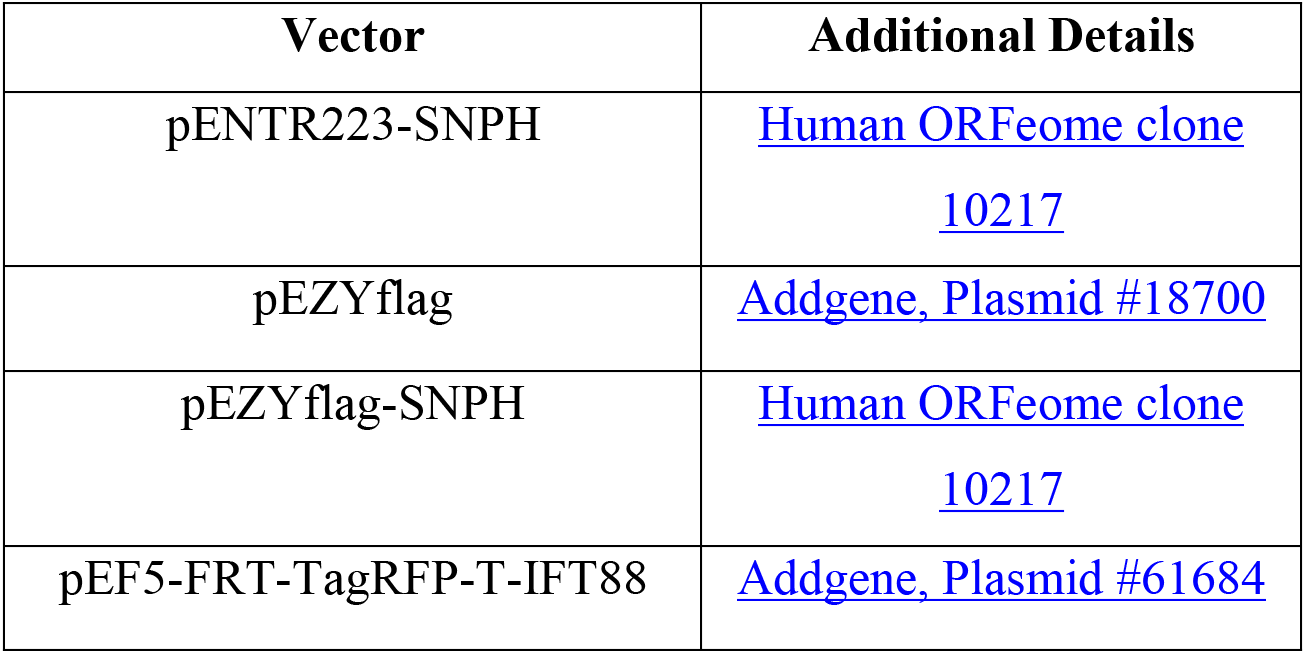

| Vector | Additional Details |
| --- | --- |
| pENTR223-SNPH | Human ORFeome clone 10217 |
| pEZYflag | Addgene, Plasmid #18700 |
| pEZYflag-SNPH | Human ORFeome clone 10217 |
| pEF5-FRT-TagRFP-T-IFT88 | Addgene, Plasmid #61684 |

#### Cloning

SNPH: The entry vector containing an *attL*-flanked gene insert encoding full-length Syntaphilin/SNPH (pENTR223-SNPH, Human orfeome clone 10217) was gateway cloned into an *attR-*destination vector encoding an N-terminal FLAG tag (Addgene, Plasmid #18700), with the Gateway™ LR Clonase™ II Enzyme mix (ThermoFisher, 11791020). DH5*<ι>α </i>*competent cells were transformed with the gateway product mix, and plated onto Carbenicllin-100 LB plates.

#### HEK293 Cell-Based Assay Autoantigen Screening

HEK293 cells were plated onto 10mm poly-d-lysine coated (50µg/mL) coverslips in 24-well plates. 293 cells were transfected overnight with pEF5-FRT-TagRFP-T-IFT88 and pEZYflag-SNPH plasmids using Lipofectamine 3000 (ThermoFisher). The following day, after two rinses with ice cold 1X PBS, FLAG-SNPH transfected cells were fixed were 4% PFA for 10 minutes, and TagRFP-IFT88 transfected cells were fixed with ice-cold methanol for 10 minutes. The fixed cells were rinsed with PBS, blocked with 5% lamb serum in PBS, and permeabilized for 30 minutes using with blocking buffer containing 0.5% Triton.

FLAG-SNPH overexpressing HEK293 cells were stained overnight with anti-FLAG antibody at 1:1000 and CSF at 1:10 in 5% blocking buffer. TagRFP-IFT88 HEK293 overexpressing cells were stained overnight using mouse anti-RFP at 1:100, rabbit anti-IFT88 at 1:100, and undiluted CSF. The cells were rinsed with PBS four times, and stained with Alexa Fluor secondaries at a 1:1000 dilution in 5% blocking buffer, FLAG-SNPH cells with anti-mouse 488 and anti-human 594, TagRFP-IFT88 cells with anti-human 488, anti-mouse 594, and anti-rabbit Cy5. Nuclei were stained with DAPI at 1:2000 in PBS for 5 minutes. Stained slides were then mounted onto microscope slides with Prolong Gold antifade.

|  | FLAG-SNPH transfected<br>HEK293 cells | TagRFP-IFT88 transfected<br>HEK293 cells |
| --- | --- | --- |
| Fixative | 2-4% PFA | Ice-cold methanol |
| Primary Stain (in 5%<br>blocking buffer) | • Rabbit anti-FLAG D6W5B (1:1000, Cell Signaling Technology (#14793))<br>• Diluted CSF (1:10) | • Mouse anti-RFP (1:100, Abcam AB65856)<br>• Rabbit anti-IFT88 (1:100, Proteintech 13967-1-AP)<br>• Undiluted CSF |
| Secondary Stain (in 5%<br>blocking buffer) | • Anti-Mouse Alexa Fluor 488 (1:1000)<br>• Anti-Human Alexa Fluor 594 (1:1000) | • Anti-Human Alexa Fluor 488 (1:1000)<br>• Anti-Mouse Alexa Fluor 594 (1:1000)<br>• Anti-Rabbit Cy5 (1:1000) |

#### Imaging

Slides were imaged at 60X at the UCSF Nikon Imaging Center using a Nikon CSU-W1 spinning disk confocal microscope, equipped with an Andor Zyla sCMOS camera.

### Immunoprecipitation-Mass Spectrometry

#### Sample preparation

For IP-MS, all sample handling through protein digestion was performed in a BSL2 biosafety hood. Technical replicates were performed on the same day, by different individuals, using different tissue preps. Plasma samples were unbuffered. CSF samples were stored 1:1 in storage buffer. Samples were thawed on ice. Immunoprecipitations were performed in unblocked, thin-walled 96-well Hard-Shell® PCR plates (Bio-Rad Cat. No. HSP9641). For immunoprecipitations using patient plasma as the source of IgG, 2 μL of plasma was diluted in 200 μL of 1x PBS (Gibco Cat. No. 10010-023). For CSF immunoprecipitations, 75 – 200 μL of CSF (depending on the amount of remaining biospecimen) was added to individual wells. For IgG conjugation, magnetic protein A/G beads were washed once in 1x PBS and suspended in an equal volume of PBS. To each well, 10 μL of washed protein A/G bead slurry was added and plates were placed on a rocker at 4C for one hour followed by incubation on a shaker at room temperature for one hour.

Postnatal day 40 – 60 mice (F1 generation of FVB x C57blk6 cross) were used as the source of antigen. Mice anesthetized in isofluorane and sacrificed by cervical dislocation. For each set, 3 brains (two males and one female for one replicate and two females and one male for the other replicate) were rapidly dissected in ice cold PBS. For each replicate, 3 brains were homogenized in ice cold tissue lysis buffer (7 mL) using a dounce homogenizer (approximately 20 strokes). Homogenized brain lysate was transferred to 1.5 mL microcentrifuge tubes and centrifuged at 4C for 10 minutes at 10,000 rcf. The supernatant from each set of brains was pooled yielding two separately prepared stocks of brain lysate. After BCA protein concentration determination, brain lysate stocks were diluted to 5 ug / uL in lysis buffer.

After IgG conjugation, the plate was placed on a magnetic plate, and the supernatant aspirated and discarded into 10% bleach. To each well, 200 μL of brain lysate (5 μg / μL) was added. Plates were sealed with adhesive aluminumized plate covers (Bio-Rad, Microseal® ‘F’ PCR Plate Seal, foil, pierceable Cat. No. #MSF1001). Antibody-bead-lysate complexes were incubated for 1 hour at room temperature under constant gentle agitation.

After 1 hour, IP plates were placed on magnetic plates and the lysate was aspirated and discarded into 10% bleach. Beads and their respective immune complexes were washed with 180 μL of 2x with detergent wash buffer, then 1x in high salt wash buffer, then 1x in nondetergent wash buffer, followed by 1x in ammonium bicarb buffer. For each well, washed beads were then resuspended in 35 μL of ammonium bicarb buffer to which 1 μL sequencing grade porcine trypsin was added (Promega, Cat. No. V5111). Immune complexes were digested on bead for 1 hr at 37°C. After digestion, plates were placed on magnetic plates and digestion reaction containing trypsinized peptides was transferred to a protein LoBind Eppendorf tube (Eppendorf, Cat. No. 022431081 and stored at −80°C until liquid chromatography (LC).

#### Spectrometry

LC separation was done on a Dionex nano Ultimate 3000 (Thermo Scientific) with a Thermo Easy-Spray source. The digested peptides were reconstituted in 2% acetonitrile/0.1% trifluoroacetic acid and 1ug in 5µl of each sample was loaded onto a PepMap 100Å 3U 75 um x 20 mm reverse phase trap where they were desalted online before being separated on a 100 Å 2U 50 micron x 150 mm PepMap EasySpray reverse phase column. Peptides were eluted using a 60 minute gradient of 0.1% formic acid (A) and 80% acetonitrile (B) with a flow rate of 200nL/min. The separation gradient was ran with 2% to 5% B over 1 minutes, 5% to 10% B over 7 minutes, 10% to 55% B over for 43 minutes, 55%B to 99%B over 1 minutes, a 4 minute hold at 99%B, and finally 99% B to 2%B held at 2% B for 10 minutes. Mass spectra were collected on a Fusion Lumos mass spectrometer (Thermo Fisher Scientific) in a data-dependent top speed mode with one MS precursor scan followed by MS/MS spectra for 3 seconds. A dynamic exclusion of 60 seconds was used. MS spectra were acquired with an isolation window of 1.2 Da, a resolution of 60,000 and a target of 4 × 105 ions or a maximum injection time of 50ms. MS/MS spectra were acquired with a resolution of 15K and a target of 1 × 104 ions or a maximum injection time of 35ms with maximum parallelizable time turned on. Peptide fragmentation was performed using collisionally induced dissociation (CID) with a normalized collision energy (NCE) value of 30. Unassigned charge states as well as +1 and ions >+5 were excluded from MS/MS fragmentation.

#### Data Analysis (Scaffold Proteome Software)

Tandem mass spectra were extracted by Proteome Explorer 1.4 (Thermo Scientific). Charge state deconvolution and deisotoping were not performed. All MS/MS samples were analyzed using X! Tandem (The GPM, thegpm.org; version X! Tandem Alanine (2017.2.1.4)). X! Tandem was set up to search the Uniprot Mouse reference proteome (1/2020, 258832 entries) and Uniprot SARS-CoV-2 Database (version 06/2020, 262 entries) assuming the digestion enzyme trypsin. X! Tandem was searched with a fragment ion mass tolerance of 0.40 Da and a parent ion tolerance of 20 PPM. Glu->pyro-Glu of the n-terminus, ammonia-loss of the n-terminus, gln->pyro-Glu of the n-terminus, deamidated of asparagine and glutamine, oxidation of methionine and tryptophan and dioxidation of methionine and tryptophan were specified in X! Tandem as variable modifications. Scaffold (version Scaffold_4.9.0, Proteome Software Inc., Portland, OR) was used to validate MS/MS based peptide and protein identifications. Peptide identifications were accepted if they could be established at greater than 95.0% probability by the Scaffold Local FDR algorithm. Peptide identifications were also required to exceed specific database search engine thresholds. X! Tandem identifications required at least. Protein identifications were accepted if they could be established at greater than 5.0% probability to achieve an FDR less than 5.0% and contained at least 2 identified peptides. Protein probabilities were assigned by the Protein Prophet algorithm (Nesvizhskii, Al et al Anal. Chem. 2003;75(17):4646-58). Proteins that contained similar peptides and could not be differentiated based on MS/MS analysis alone were grouped to satisfy the principles of parsimony. Proteins sharing significant peptide evidence were grouped into clusters. For identification of significant peptides, Scaffold settings were as such: protein threshold = FDR < 5%, minimum peptides = 1, peptide threshold = MRS_otIt. Mouse proteins that were significantly enriched after Benjamini-Hochberg corrected t-test (alpha = 0.05) and that were observed in both replicates were considered candidate autoantigens.

#### Data Analysis (MS1 Peak Area – Quandenser)

The Quandenser pipeline (https://github.com/statisticalbiotechnology/quandenser-pipeline) was used with its default settings on the raw mass spectrometry files. The Quandenser pipeline consists of Quandenser, Crux and Triqler (cite = https://www.nature.com/articles/s41467-020-17037-3)

#### Data Analysis (Gene ontology)

ToppGene.org was used for gene ontology analyses of IP-MS and PhIP-Seq Data.

### Phage Display Immunoprecipitation Sequencing (PhIP-Seq)

#### Preparation of phage libraries from stocks

A 500mL culture of E. Coli BLT5403 was incubated at 37°C with shaking until log phase, defined as OD_600_= 0.5. The culture was then inoculated with the stock phage library with a multiplicity of infection (MOI) of 0.001 and incubated at 37°C for 3 hours in order for complete lysis to occur. Lysate was spun at 3500rpm for 15 minutes to remove debris.

Phage were precipitated by adding 20% culture volume of 5x PEG/NaCl (PEG-8000 20%, NaCl 2.5mM) and left overnight at 4°C. Phage were then spun down at 4°C for 15 minutes at 10,000g, re-suspended in storage media (20 mM Tris-HCl, pH 7.5, 100 mM NaCl, 6mM MgCl_2_) and filtered through a 0.22uM filter. Resulting phage libraries were then tittered by plaque assay, with an acceptable working concentration between 10^9^-10^11^ pfu/mL.

#### Immunoprecipitation of phage-bound patient antibodies

96-well 250uL BioRad hardshell plates were blocked overnight with blocking buffer (3% BSA, 0.1% PBS-T) to prevent nonspecific binding of phage and/or antibodies to walls of the plate. Patient samples were stored in blocked 96-well plates and diluted in 2x storage buffer (40% glycerol, 40mM HEPES pH7.3, 0.04% NaN_3_ in DPBS). Blocking buffer was then replaced with 150uL of pre-prepared phage library and incubated with 2uL patient sample overnight with shaking at 4°C. Commercial antibodies were diluted at 1:50.

Protein A and G magnetic beads (Thermo-Fisher Scientific) were washed by magnetic separation and re-suspended in an equivalent volume of 0.1% TNP-40 (150mM NaCl, 50mM Tris-HCL, 0.1% NP-40). 10uL of A/G beads were added to each IP reaction and incubated at 4°C with shaking for 1 hour to allow patient antibodies to bind to beads. Beads were then washed by magnetic separation with 150uL 0.1% TNP-40 three times. Subsequent to final wash, beads were re-suspended in 150uL of Luria broth (LB) and used to inoculate 500uL of fresh E. Coli cultures (OD_600_= 0.5) in 96-deep well plates for in vivo amplification. Cultures were incubated for 2 hours at 37°C with shaking and allowed to complete lysis. 96-deep well plates were then spun down at 3300prm for 10 minutes. 150uL of lysate (IP1) was saved and stored at −20°C. Another 150uL of lysate was mixed with an additional 2uL of patient sample overnight 4°C and immunoprecipitation was repeated (IP2) to further enrich for relevant antibody-binding phage.

#### NGS library preparation of phage DNA

Plates containing lysate from IP2 were thawed and subsequently diluted 1:50 in ddH_2_O. Enrichment of phage DNA and barcoding of individual IP reactions was performed using a single PCR reaction using multiplexing primers and the following reagents:

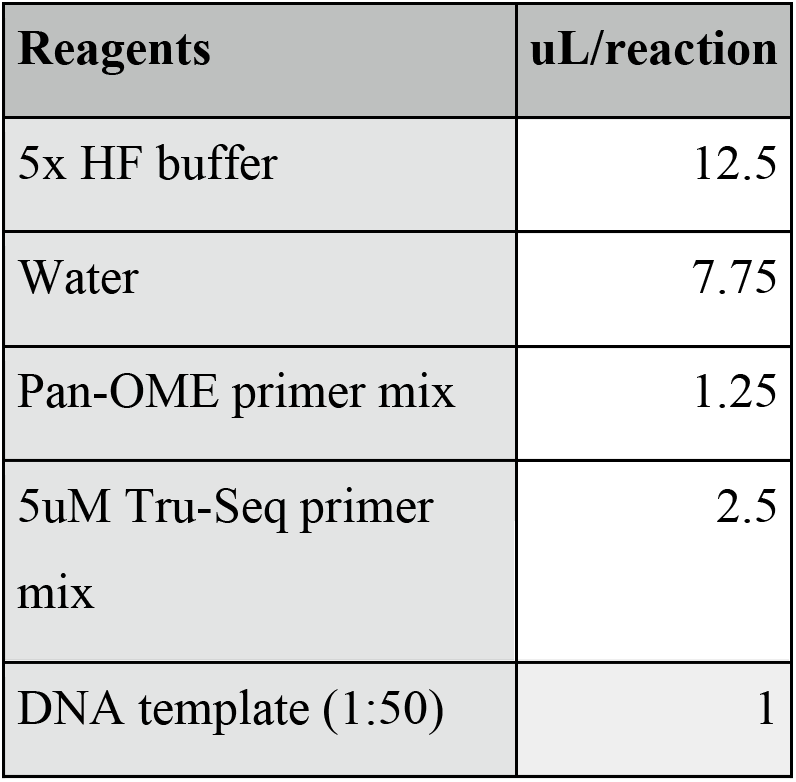

| Reagents | uL/reaction |
| --- | --- |
| 5x HF buffer | 12.5 |
| Water | 7.75 |
| Pan-OME primer mix | 1.25 |
| 5uM Tru-Seq primer mix | 2.5 |
| DNA template (1:50) | 1 |

### Primer design for pan-OME primers

panOME_rev0:

GTGACTGGAGTTCAGACGTGTGCTCTTCCGATCTTAGTTACTCGAGTGCGGCCGCAAGC

panOME_rev1:

GTGACTGGAGTTCAGACGTGTGCTCTTCCGATCTNTAGTTACTCGAGTGCGGCCGCAAGC

panOME_rev2:

GTGACTGGAGTTCAGACGTGTGCTCTTCCGATCTNNTAGTTACTCGAGTGCGGCCGCAAGC

panOME_rev3:

GTGACTGGAGTTCAGACGTGTGCTCTTCCGATCTNNNTAGTTACTCGAGTGCGGCCGCAAGC

panOME_forward_MP

ACACTCTTTCCCTACACGACGCTCTTCCGATCTAGTCAGGTGTGATGCTCGGGGATCC

### Primer design for multiplexing primers

Forward:

AATGATACGGCGACCACCGAGATCTACAC[NNNNNNN]GGAGCTGTCGTATTCCAGTCAGGTGTGATGCTC

NNNNNNN = i5 index

Reverse:

CAAGCAGAAGACGGCATACGAGAT[NNNNNNN]GGTTAACTAGTTACTCGAGTGCGGCCGCAAGC

NNNNNNN = i7 index

### Thermocycling conditions

1. 95°C for 2 minutes
2. 5 cycles of: 95°C ×30sec, 63°C ×30sec, 72°C ×45sec
3. 5 cycles of: 95°C ×30sec, 60°C ×30sec, 72°C ×45sec
4. 18 cycles of: 95°C ×30sec, 58°C ×30sec, 72°C ×45sec
5. 72°C for 2 minutes
6. hold at 10°C

After PCR reaction was completed, products were immediately pooled (3uL per sample) and size selected with a 0.67x bead clean-up (SPRISelect, Beckman Coulter). Product concentration was quantified using an automated electrophoresis system (Bioanalyzer, High-Sensitivity DNA Chips, Agilent). Libraries were sequenced on the iSeq (cases 1 - 6 and controls) or NovaSeq 6000 (case 7 and controls) (Illumina) using 150nt paired-end reads.

#### PhIP-Seq Bioinformatic Analysis

Raw reads generated from the PhIP-Seq peptidome assay were aligned to the reference database using RAPSearch (v2.2). Peptide counts outputted from this workflow were normalized to reads per 100 thousand (RPK) for every sample by diving each peptide count by the sum and multiplying by 100,000. The resulting peptide RPK count matrices were analyzed using a custom bioinformatics pipeline written in R.

For the analysis, these data were divided into disease and reference groups for both cerebrospinal fluid (CSF) and plasma samples. The disease group contained SARS-CoV-2 and A-G bead samples. The reference group contained healthy control (HC) and A-G bead samples. Peptide Fold Change (FC) was calculated for each sample. Peptide counts for Sars-CoV2 samples were divided by the mean RPK of the reference group and healthy control samples were divided by the mean RPK of the disease group. In addition, the FC for GFAP samples were calculated in the same way using the mean of all A-G bead samples.

To identify enriched peptides, results from each sample were filtered using a set of thresholds that, when using a commercial antibody to GFAP, consistently identified GFAP peptides while minimizing nonspecific off-target peptide identification (Agilent Z033429-2, 1:200 dilution). Each peptide was required to have a minimum of 1 RPK as well as a FC > 10. In addition, thresholds were applied at the gene level. Genes were kept if at least one peptide had a FC > 100 and a total (summed) RPK > 20 across all peptides in the gene. A Kmer analysis was applied to amino acid sequences of all peptides that passed the previous filters. Using a sliding window algorithm, with a window size of 7 and a step size of 1, all 7-mers were compared across SARS-CoV-2 and HC samples. Proteins for which peptides containing at least one 7-mer overlap with another peptide whose total rpK was ≥ 20 were carried forward in the analysis.

Additionally, proteins with nonoverlapping peptides with an individual rpK ≥ 20 an FC ≥ 100 were also carried forward. Proteins that passed these thresholds in both technical replicates but were not enriched by reference samples were considered candidate autoantibodies. This workflow was repeated on a per sample level, and the results for each sample were stored separately.

### Human Monoclonal antibodies

#### Expression Vector Cloning

Select heavy and light chain variable region fragments, or framework 1 through 4 as defined by the IMGT human V gene database, were synthesized by Integrated DNA Technologies, Inc. Heavy, kappa, and lambda fragments were cloned into AbVec2.0-IGHG1 (Addgene plasmid # 80795; http://n2t.net/addgene:80795; RRID:Addgene_80795), AbVec1.1-IGKC (Addgene plasmid # 80796; http://n2t.net/addgene:80796; RRID:Addgene_80796), and AbVec1.1-IGLC2-XhoI (Addgene plasmid # 99575; http://n2t.net/addgene:99575; RRID:Addgene_99575) linearized expression vectors, respectively. All expression vectors were a gift from Hedda Wardeman(*50*). Cloning was performed in a total of 20ul with 60ng of linearized vectors, 18ng heavy or light chain fragment and 10ul of Gibson assembly master mix (New England Biolabs). 5-alpha competent *E. coli* (New England Biolabs) were transfected at 42C with 2ul unpurified, assembled plasmid. Colonies were sequence to confirm correct assembly, and then plasmids were purified from 3mL of *E. coli* in LB Broth with 100ug/mL ampicillin using Qiaprep Spin columns (Qiagen).

#### Monoclonal Antibody Production

Monoclonal antibodies (mAbs) were produced by Celltheon Co. by transfecting Chinese hamster ovarian cells with equal amounts of heavy chain and light chain plasmids. Culture supernatant was harvested and mAbs were subjected to affinity purification.

#### Monoclonal autoantibody screening

Human-derived monoclonal antibodies were screened by Luminex serology (1 ug per replicate), anatomic mouse brain tissue staining (18ug/mL), PhIP-Seq (0.06ug mAb per immunoprecipitation, 2 immunoprecipitations per replicate), and IP-MS (5 ug per IP).

### Data availability

Gene expression and repertoire data in the study are available in the NCBI repository SRAxxxx Raw mass spectrometry files and data analysis output are available from MassIVE (https://massive.ucsd.edu/) and Proteome Exchange (http://www.proteomexchange.org/) under MassIVE dataset identifier # and Proteome Exchange #

### Schematics

Schematic images were created with Biorender.com

## Supplementary Materials

**Fig. s1.**
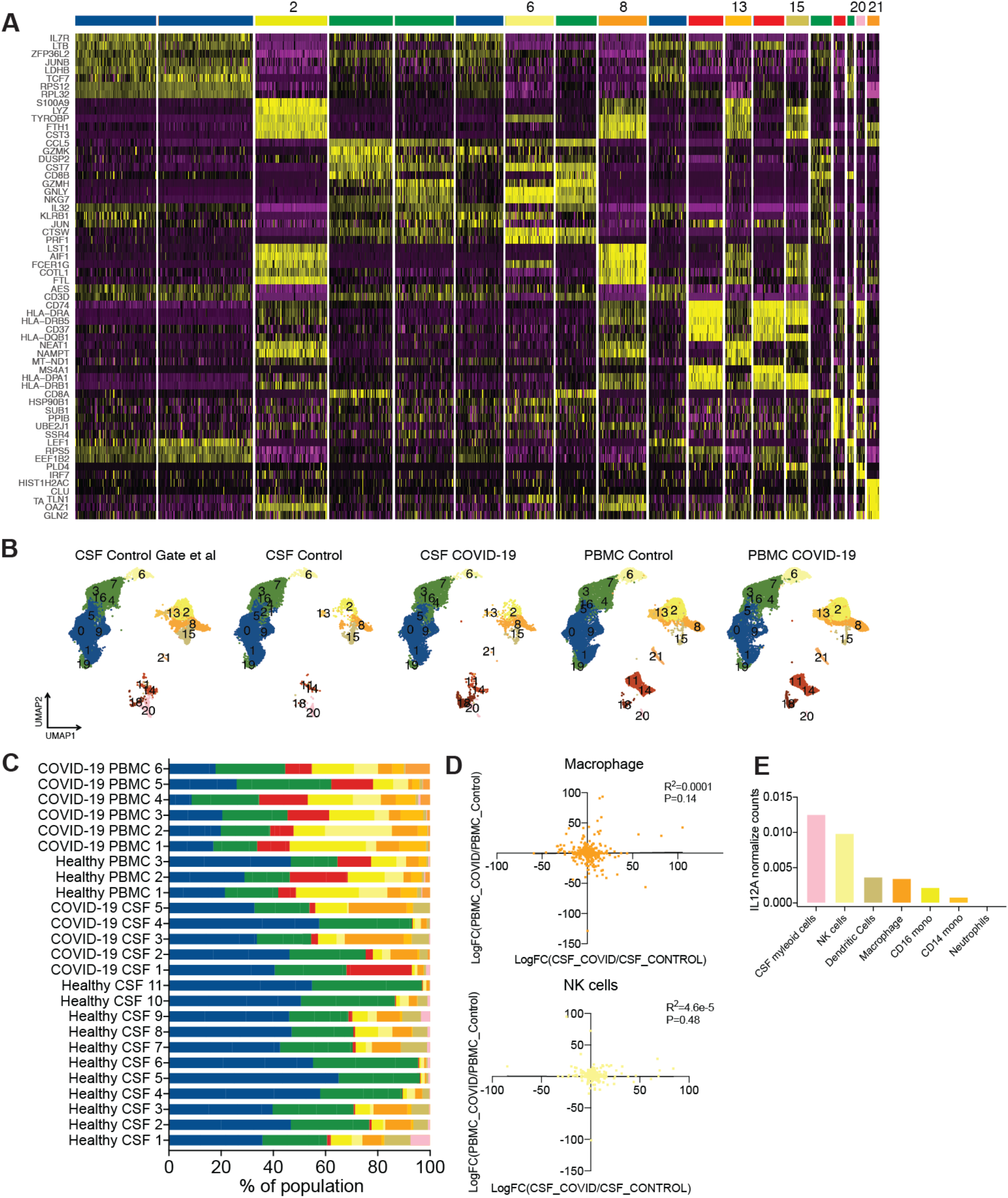
Distinct immunological landscape of COVID-19 patient’s CSF versus PBMC. (**A**) Top DEGs distinguishing clusters found in Figure 1. (**B**) UMAP projection of cell types in CSF and PBMC of COVID-19 cases and healthy controls. (**C**) Relative proportion of cell types found in biological samples for the study. (**D**) Correlation of Log fold change of genes in COVID-19 patient CSF versus PBMC in macrophages and NK cells. (**E**) Normalized counts of IL12A transcripts on a population level for innate immune cells.

**Fig. s2:**
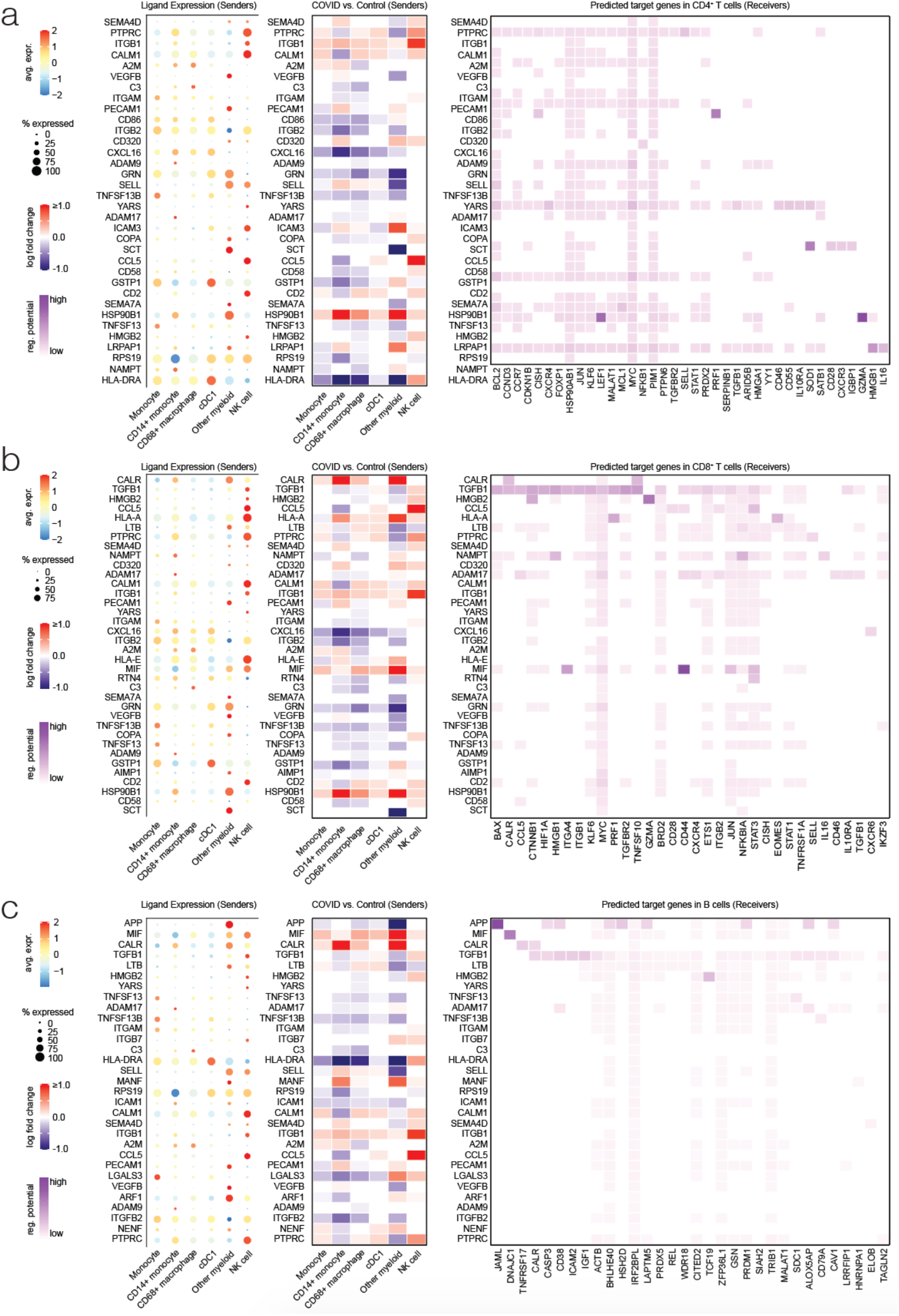
Interaction of innate and adaptive immune cells predicted by NicheNet. NicheNet analysis of innate immune cell interaction with CD8 T cells. Left panel shows relative expression levels of ligands expressed in different innate immune cell compartments. Middle panel shows the change in each ligand in COVID-19 patient CSF versus control patient CSF. Right panel shows a heatmap of probability of regulatory potential of these ligands (y-axis) to affect the differentially expressed genes in the receiver. (**A**) Shows analysis with receivers as CD4 T cells. (**B**) Shows analysis with receivers as CD8 T cells. (**C**) Shows analysis with receivers as B cells.

**Fig. s3.**
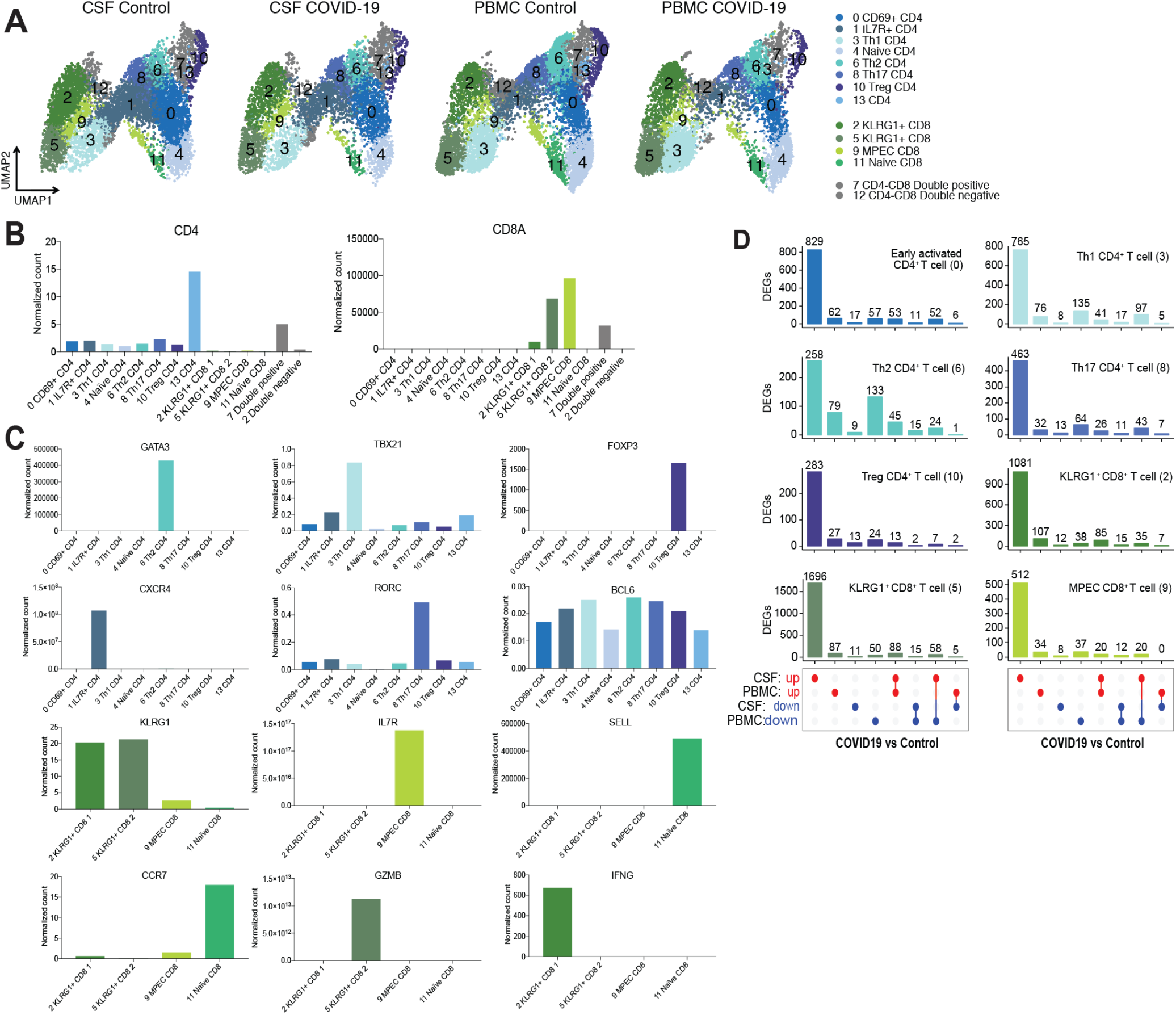
Characterization of T cell subsets found in CSF and PBMC. (A) UMAP projection of T cell types in CSF and PBMC of COVID-19 patients and control patients. (**B, C**) Normalized gene expression data of T cell identification genes in T cells. **(D)** UpSet plot showing differentially expressed genes (DEGs) in T cell subsets of COVID-19 cases versus healthy controls in CSF and PBMC. Each column denotes the number of genes that are up or down-regulated in CSF and/or PBMC as indicated in the bottom panel.

**Fig. s4:**
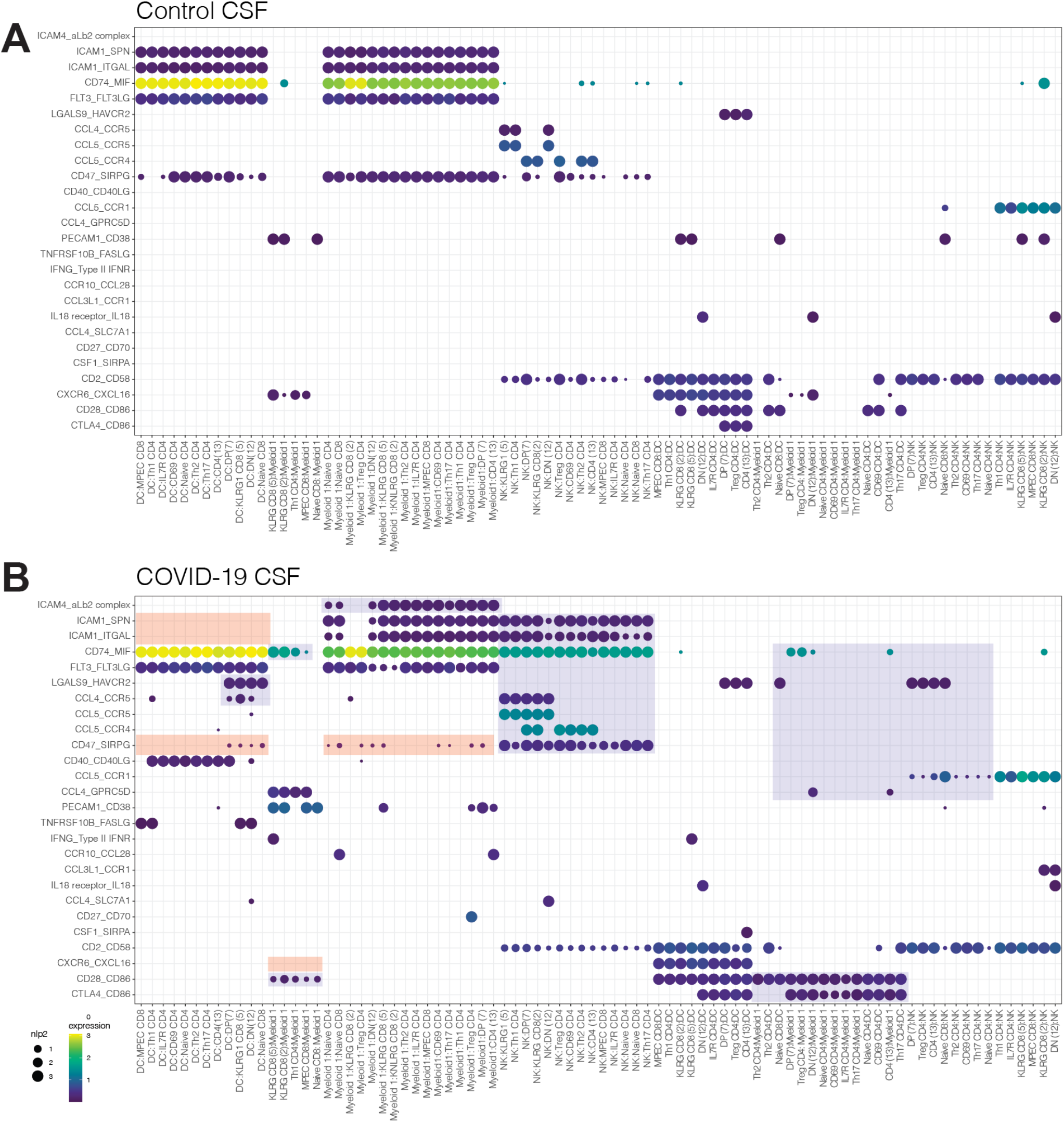
Predicted interaction enriched in CSF of COVID-19 patients using CellPhoneDB. CellPhoneDB interaction map between innate immune cells identified in Fig 1A and T cell subtypes identified in Fig 2A of COVID-19 patient CSF. (**A**) Depicts control patient CSF interactions. (**B**) Depicts interactions in CSF of COVID-19 cases. Red box depicts interactions disappearing compared to control CSF and blue box depicts interactions enriched compared to control CSF.

**Fig s5:**
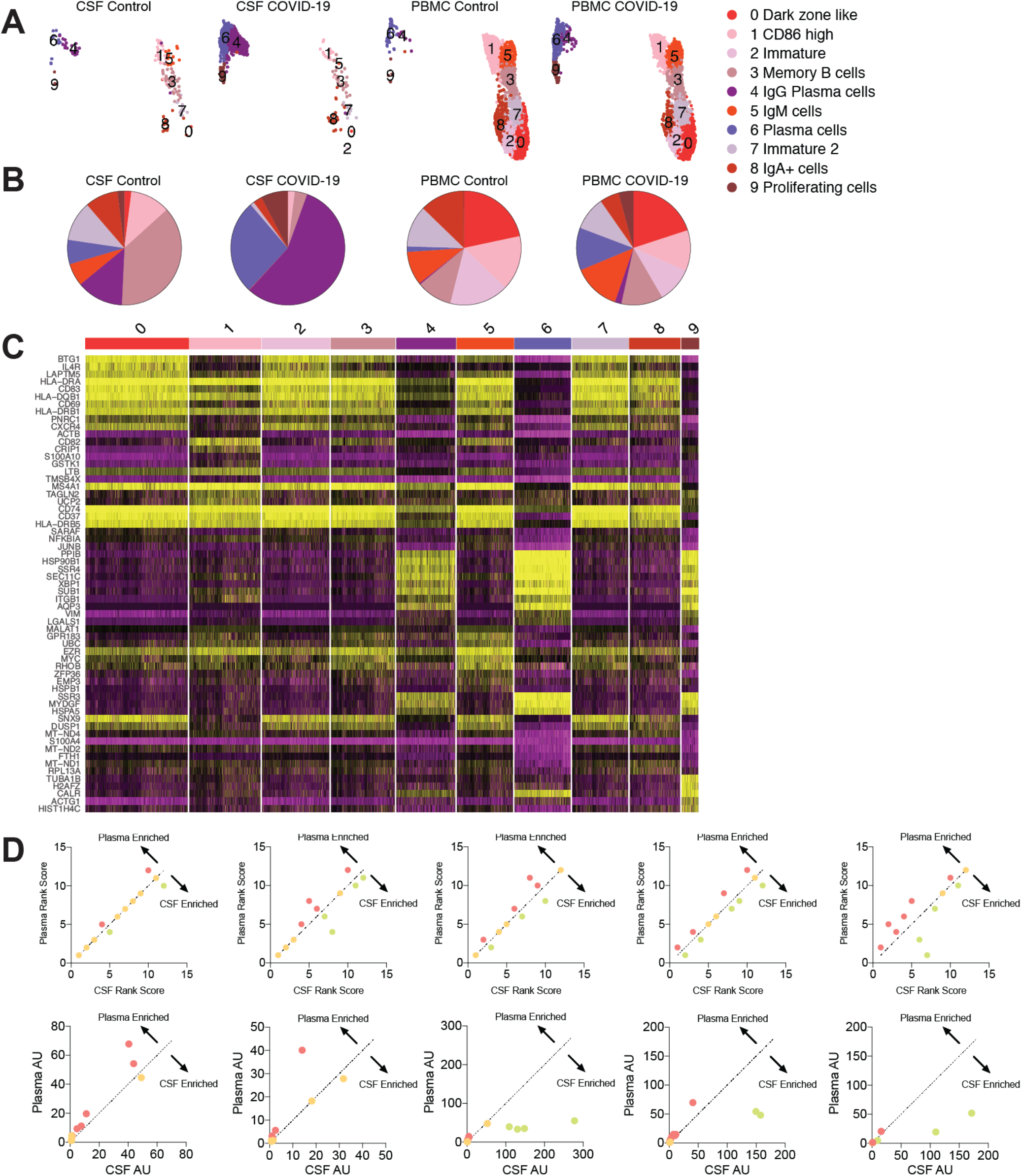
CNS localized B cell responses occur in COVID-19 patients. (**A**) UMAP projection of B cell types in CSF and PBMC of COVID-19 cases and controls. (**B**) Pie charts depicting relative population frequency of different B cell subtypes found in CSF and PBMC of COVID-19 cases and controls. (**C**) Top DEGs distinguishing clusters found in Figure 3. D**d, top row**) SARS-CoV-2 epitope binding antibody frequency from Figure 3c was ranked and correlation between CSF and plasma was performed (higher rank score depicts more frequent epitope. (**D, bottom row**) Similar to top row, but normalized arbitrary unit was used to derive a correlation between plasma and CSF antibodies. Each panel represents individual patient samples. Each dot represents different epitopes. Green represents epitopes enriched in CSF, Red represents epitopes enriched in Plasma, Orange represents epitopes that are not enriched in either compartment.

**Fig s6:**
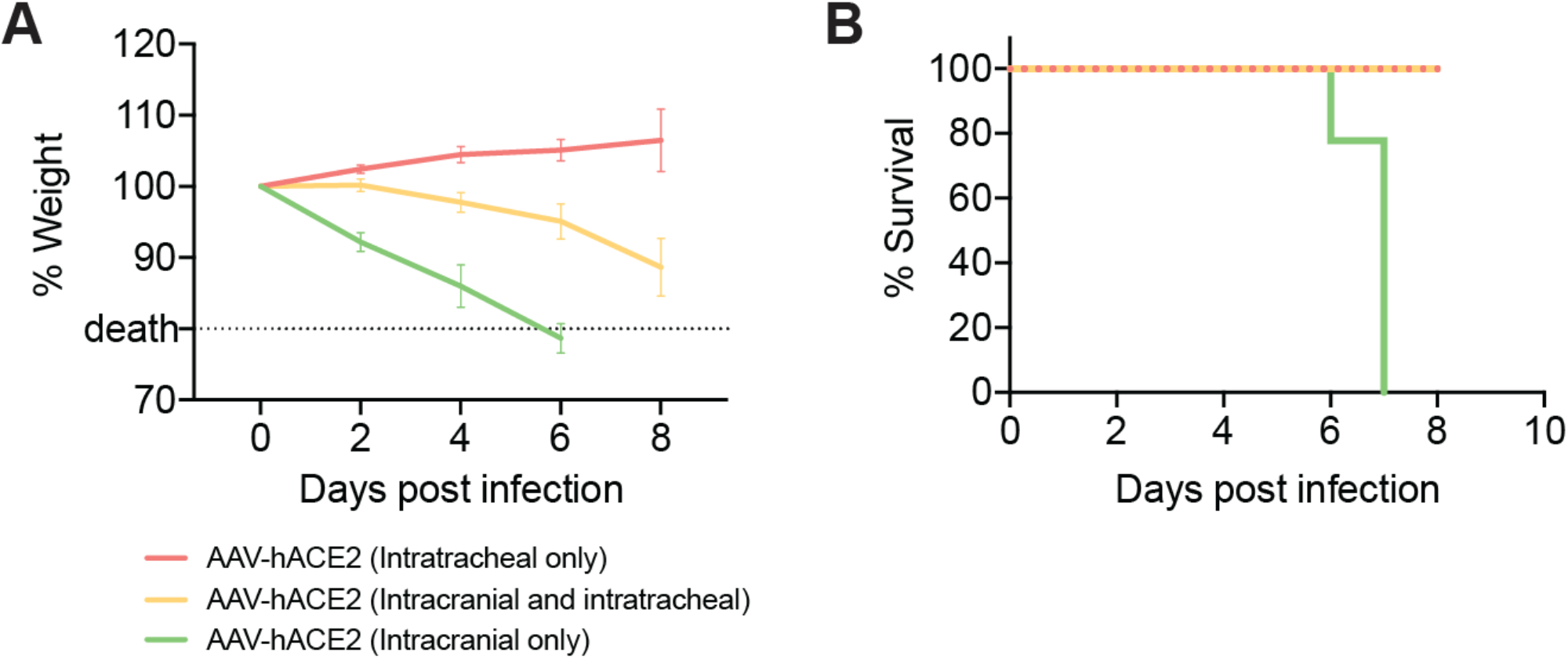
Weight loss and survival of mice infected with SARS-CoV-2. Refer to Figure 4 for experiment design. (**a**) Weight loss curve for mice described in Figure 4. (b) Kaplan meier curve of mice described in Figure 4 (death was determined by weight loss).

**Figure s7.**
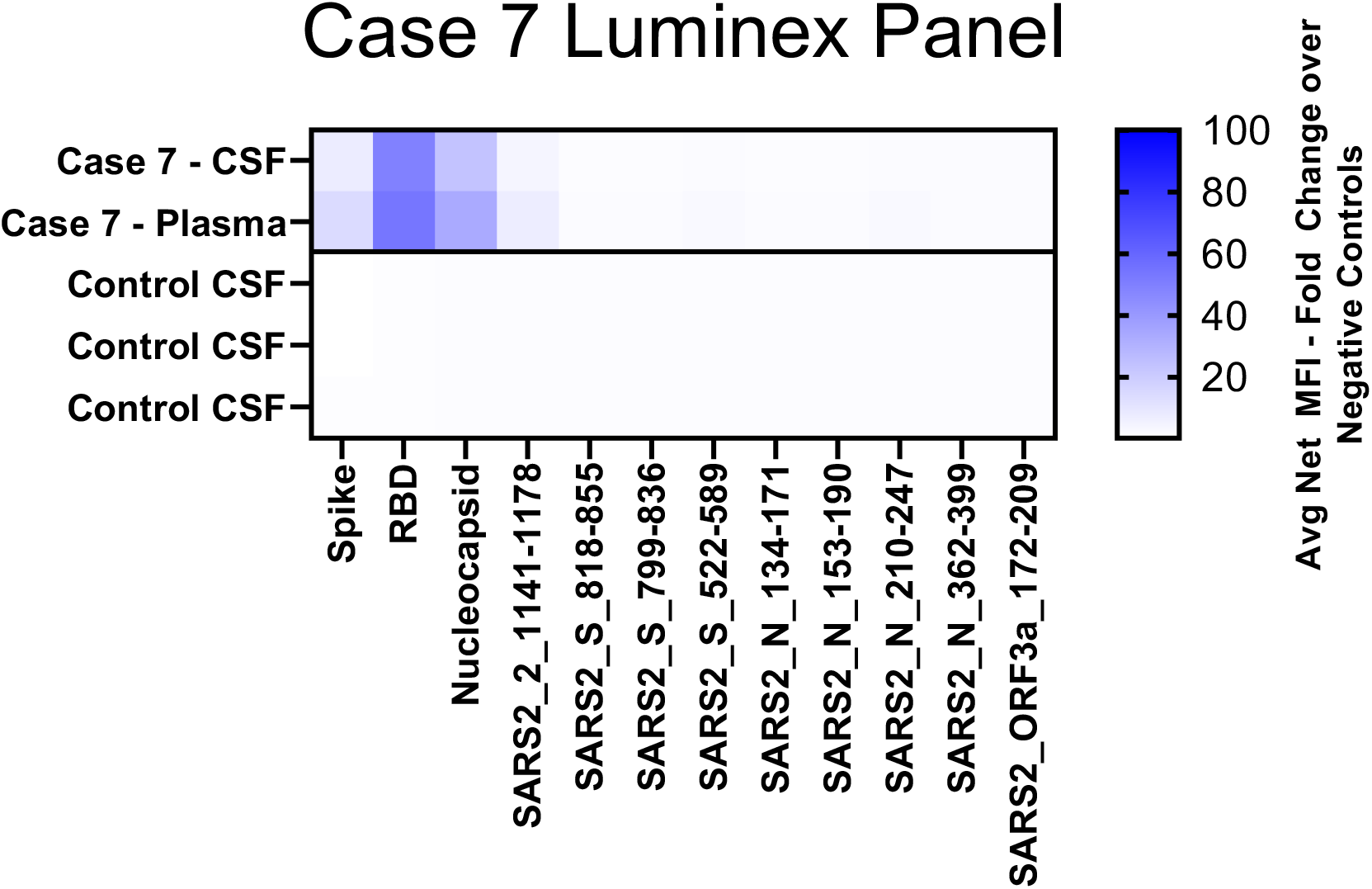
Luminex panel demonstrating SARS-CoV-2 antibodies in post-COVID-19 case 7. Heatmap of anti-SARS-CoV-2 antibody specificity of CSF and plasma of case 7.

**Fig s8.**
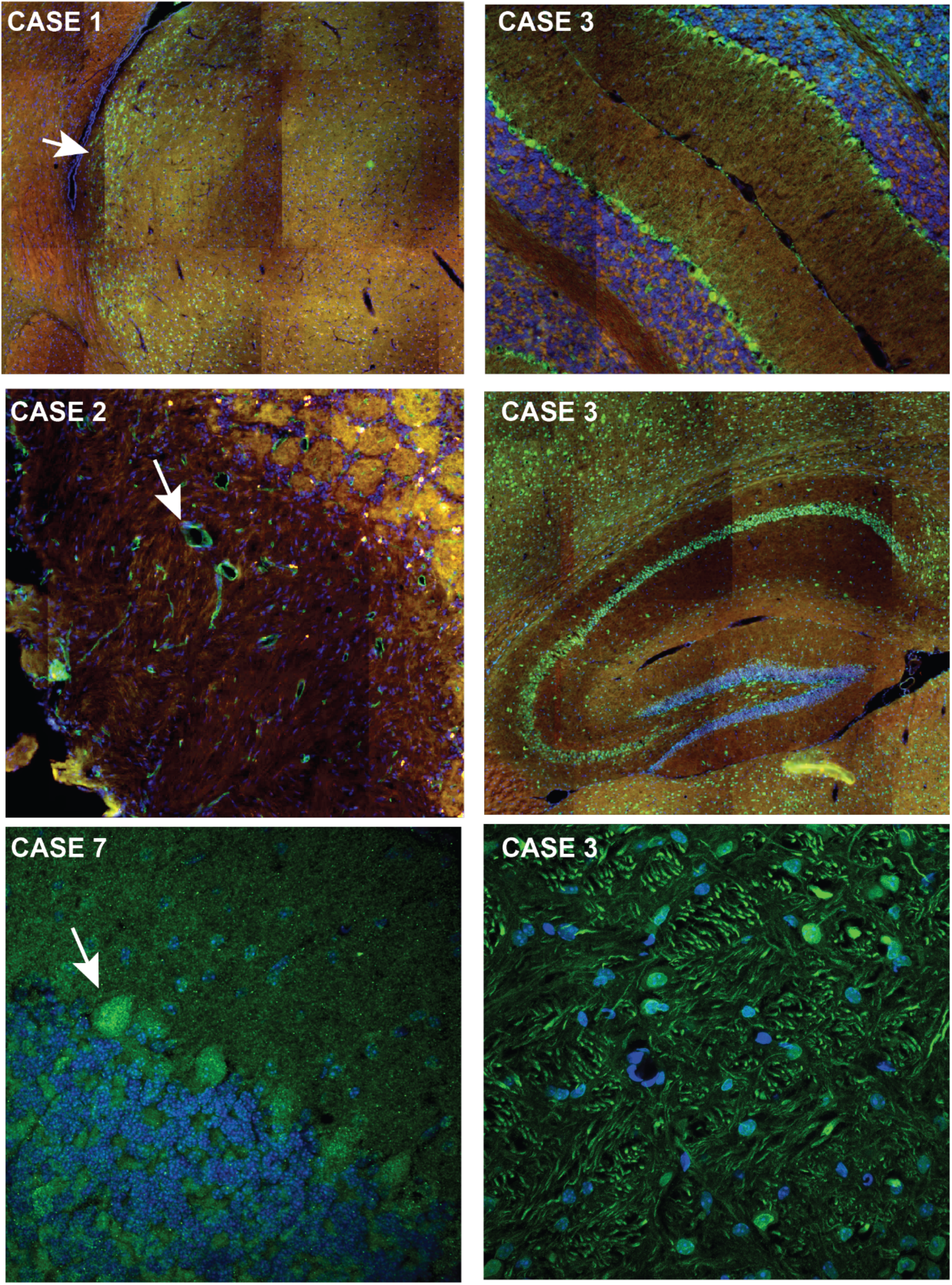
Additional anatomic regions immunostained by COVID-19 CSF at a 1:10 dilution. Clockwise from top left: (1) Case 1, anterior paraventricular nucleus of the thalamus, (2) Case 3, Purkinje cells and glial like processes in the cerebellum, (3) Case 3, hilus, CA fields of the hippocampus, subiculum, and overlying cortical cells, (4) Case 3, anterior pons of the brain stem, (5) Case 7, Purkinje cell bodies and synaptic-like puncta in the overlying molecular layer of the cerebellum, (6) Case 2, endothelial-like cells in the olfactory bulb.

**Fig s9.**
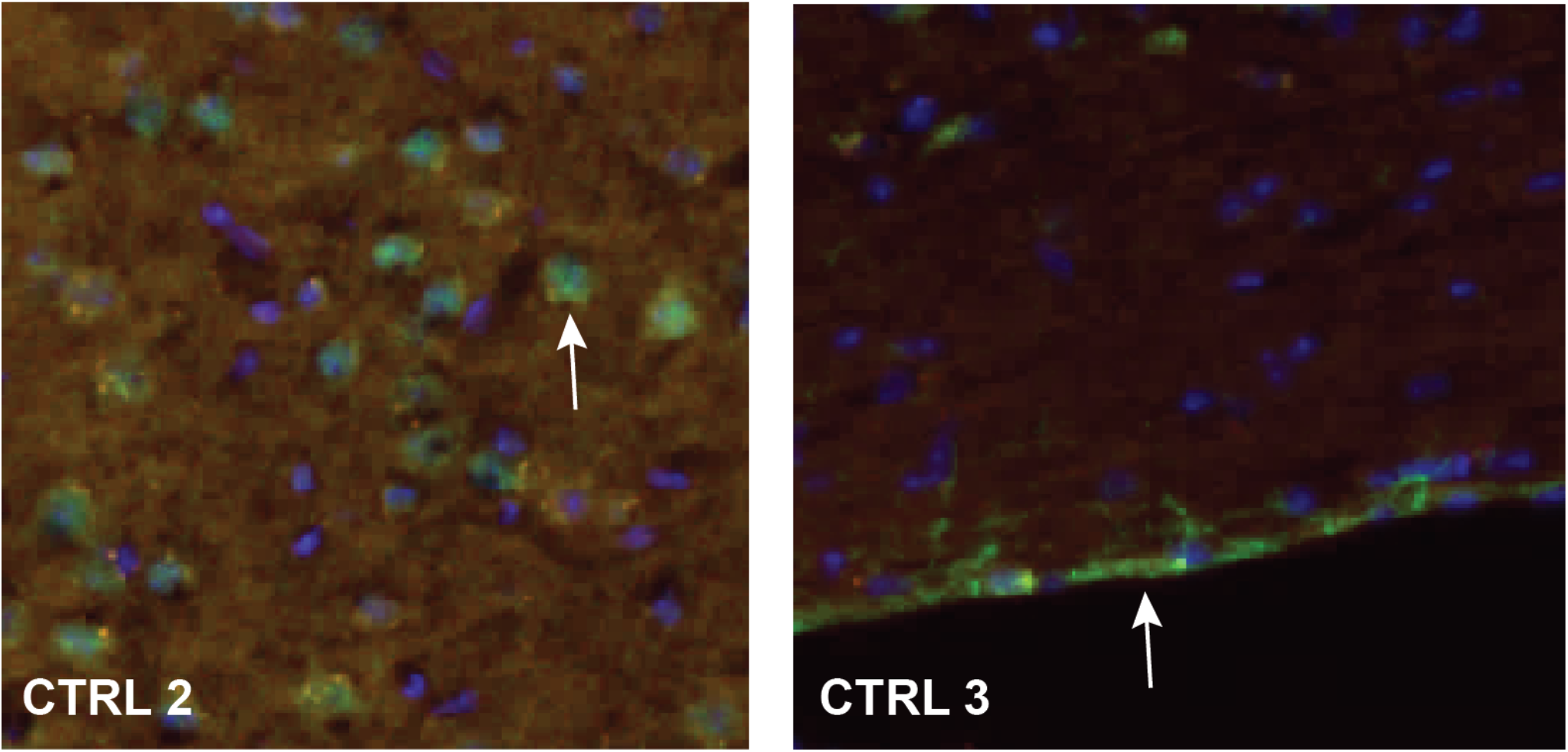
Control CSF immunoreactivity. Two of six control CSF samples were immunoreactive to mouse brain tissue at a 1:10 dilution but not at higher dilutions. CTRL 2 (left) was weakly nuclear throughout the brain. CTRL 3 was primarily immunoreactive to subpial glial.

**Fig s10.**
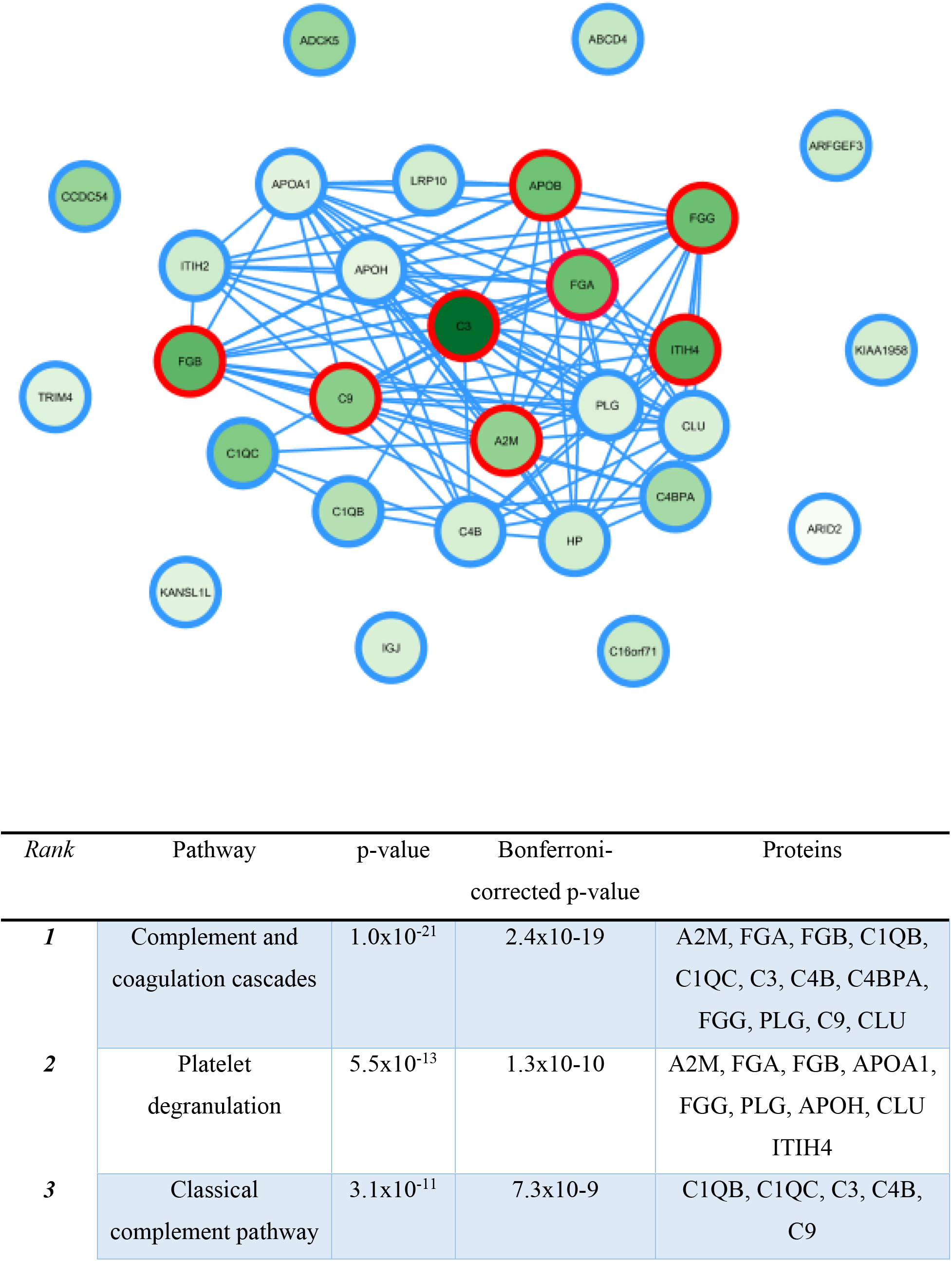
Network analysis and gene ontology pathway analysis of IP-MS identified human plasma proteins from COVID-19 patients. Top network: COVID-19 plasma immunoprecipitations were compared as a group to control plasma immunoprecipitations. Human proteins that were elevated 1.5-fold or greater by MS1 peak areawere analyzed by string-db.org. String-db network connection data were imported into cytoscape along with IP-MS q-values (a q-vlaue is an FDR-adjusted p-value). The protein nodes are shaded (green) according to their log2 fold enrichment. Red borders indicate those proteins with a q-value < 0.05, and blue borders indicate q-values > 0.05. Bottom table: ToppGene gene ontology pathway anaylsis of the set of proteins (28/29, IGJ was excluded) that were elevated 1.5-fold or greater in the plasma IgG IP-MS fractions of COVID-19 patients.

**Fig s11.**
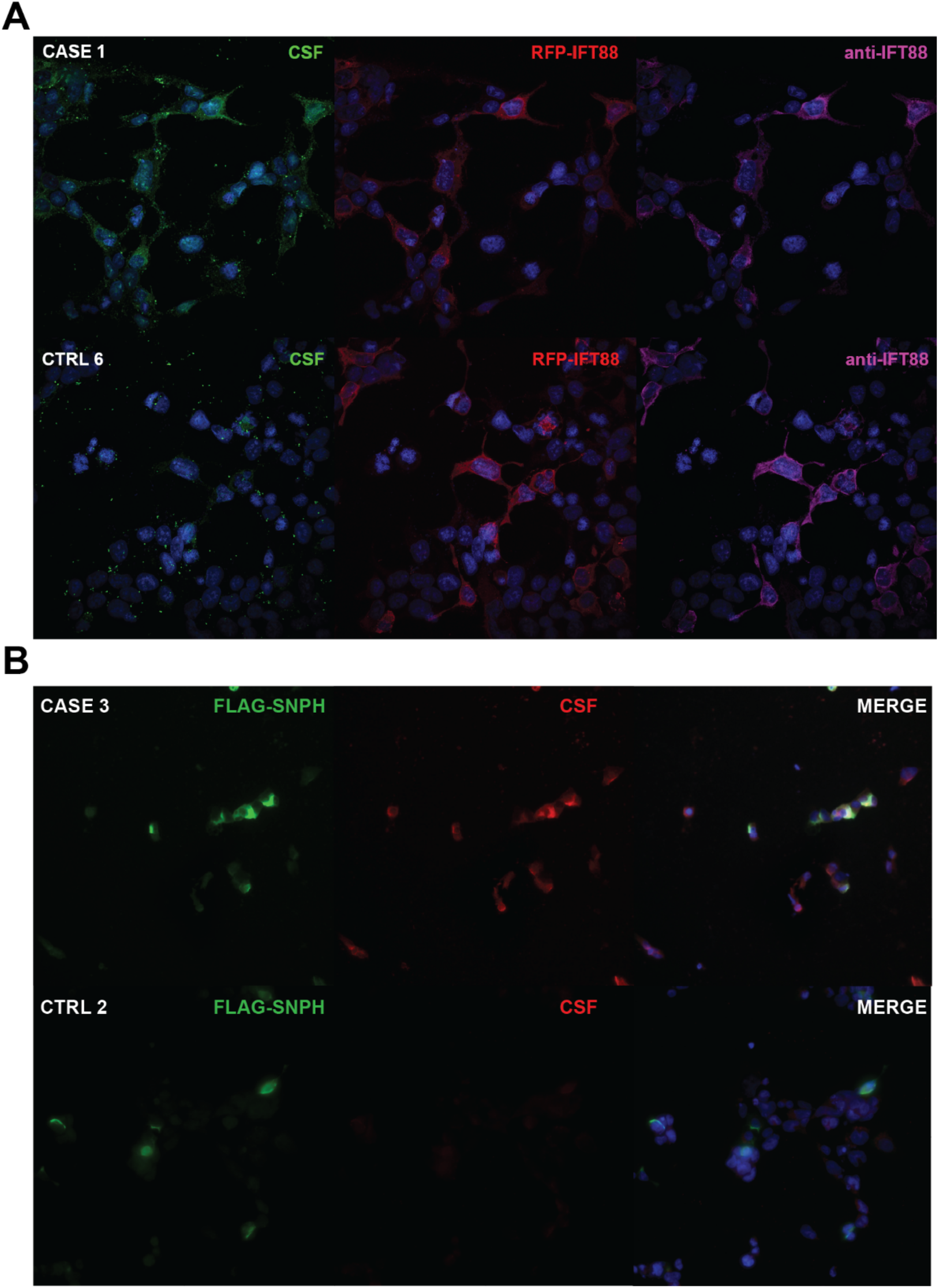
HEK 293 overexpression cell based assay for IFT88. **(A)** Confocal images of IFT-88 CBA. Top row: Case 1 CSF (green) immunoreacts to overexpressed RFP-IFT88 (red) and colocalizes with a commercial anti-IFT88 antibody (magenta). Bottom row: Control CSF (green) is not immunoreactive RFP-IFT88 (red) and does not colocalize with a commercial anti-IFT88 antibody (magenta). **(B)** Nikon images of SNPH CBA. Top Row: Case 3 CSF (red) is immunoreactive to overexpressed FLAG-SNPH (green) and colocalizes (merge). Bottom row: Control 2 CSF (red) is not immunoreactive to overexpressed FLAG-SNPH (green).

**Table S1.**
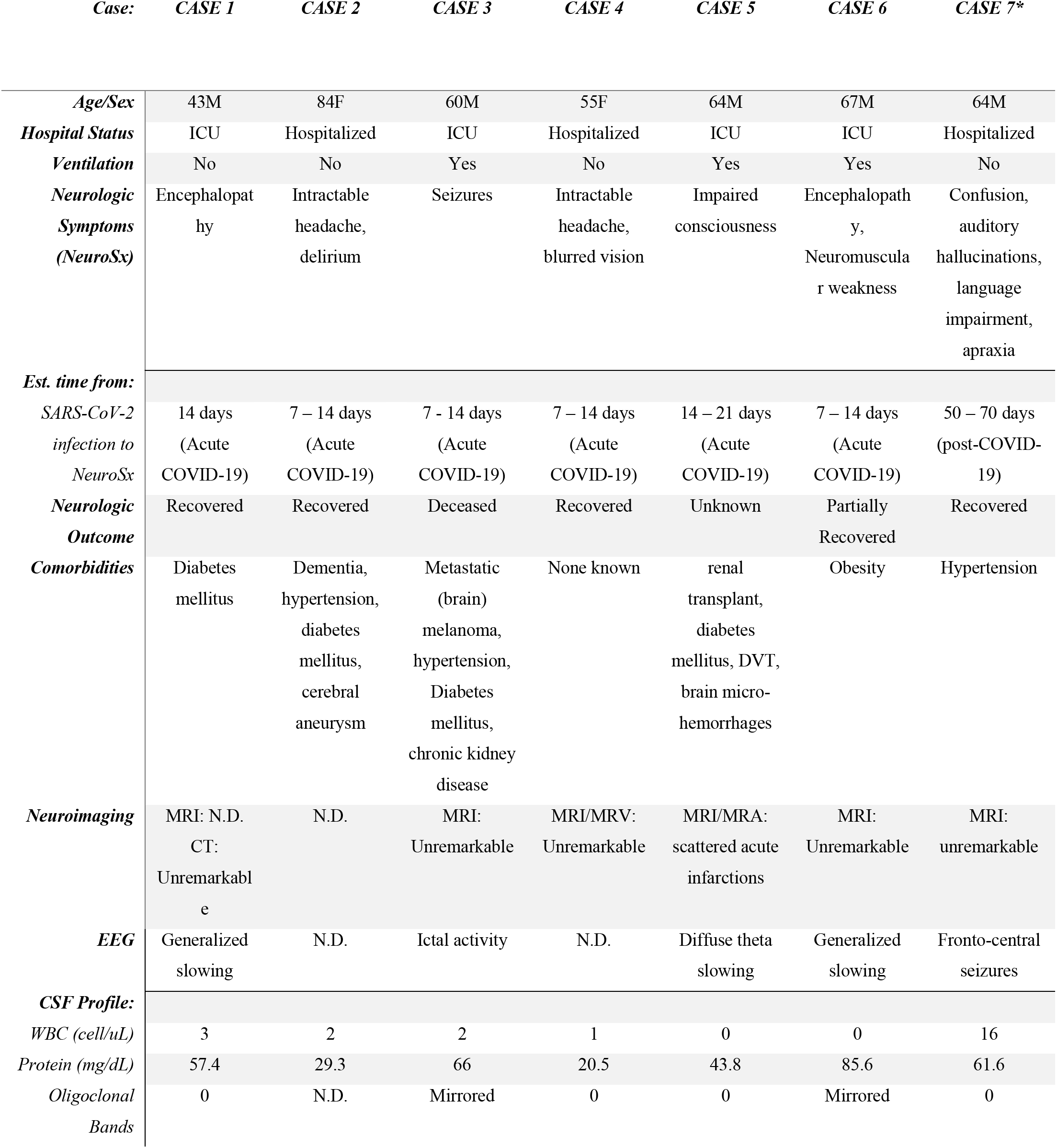
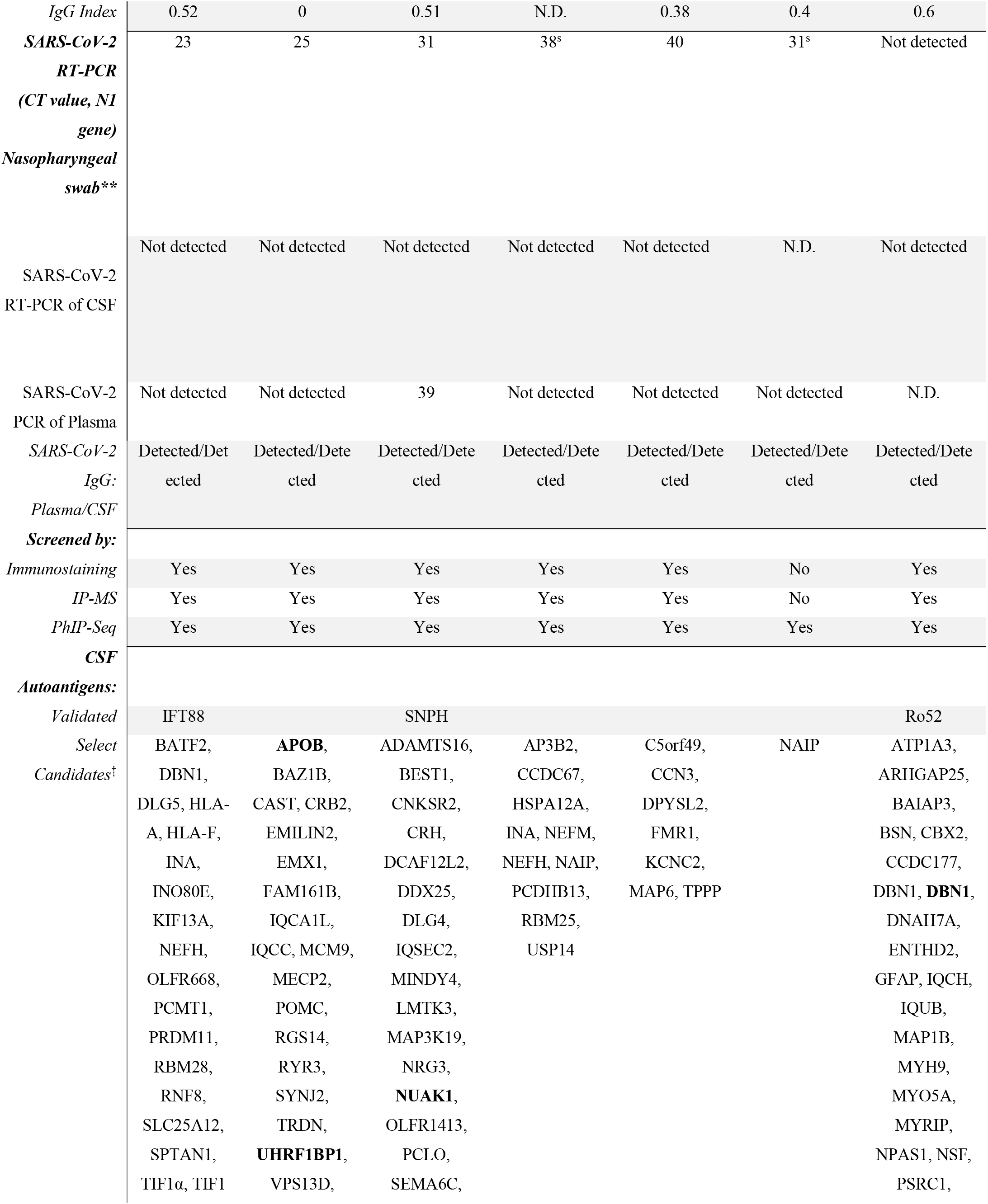
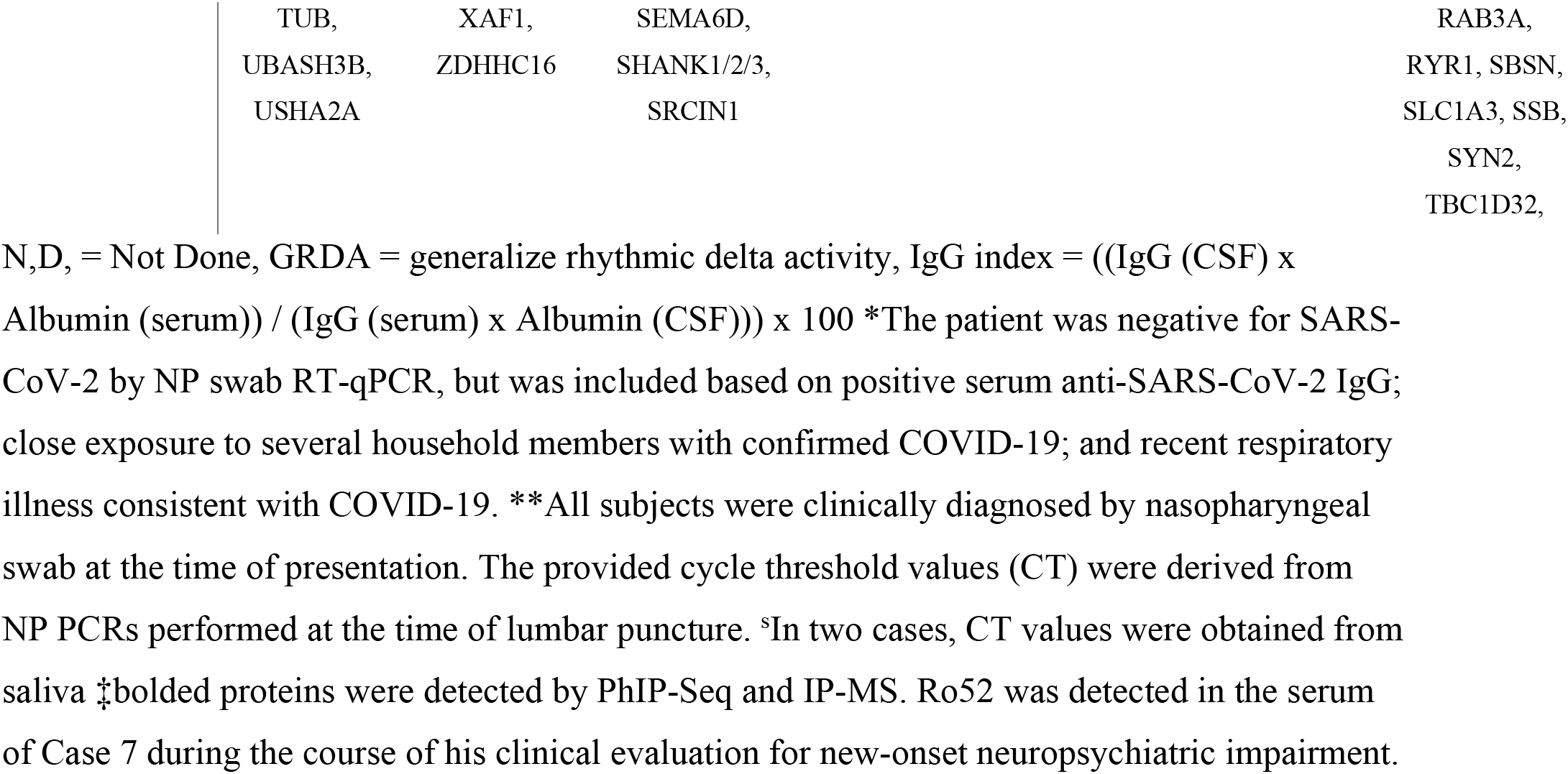
COVID-19 Patient Characteristics.

**Table S2.** Control Characteristic.

| *Case:* | *CTRL 1* | *CTRL 2* | *CTRL 3* | *CTRL 4* | *CTRL 5* | *CTRL 6* | *CTRL 7\** | *CTRL 8\** | *CTRL 9\** | *CTRL 10\** | *CTRL 11\** | *CTRL 12\** | *CTRL 13\** |
| --- | --- | --- | --- | --- | --- | --- | --- | --- | --- | --- | --- | --- | --- |
| *Age/Sex* | 75M | 54M | 54M | 58M | 45F | 49M | 23M | 32F | 49M | 52M | 33M | 52M | 56M |
| *Neurologic Symptoms* | Lower Extremity polyneuropathy | None | None | None | None | None | None | None | None | None | None | None | None |
| *Comorbidities* | Alcohol use disorder | None Known | None Known | None Known | None Known | None Known | None Known | None Known | None Known | None Known | None Known | None Known | None Known |
| *CSF Profile:* |  |  |  |  |  |  |  |  |  |  |  |  |  |
| *WBC (cell/uL)* | 12 | 2 | 1 | 0 | 3 | 1 | N,D, | N,D, | N,D, | N,D, | N,D, | N,D, | N,D, |
| *Protein (mg/dL)* | 40,7 | 37 | 48,7 | 43,5 | 38,3 | N,D, | N,D, | N,D, | N,D, | N,D, | N,D, | N,D, | N,D, |
| *OCBs* | 0 | N,D, | N,D, | N,D, | N,D, | N,D, | N,D, | N,D, | N,D, | N,D, | N,D, | N,D, | N,D, |
| *SARS-CoV-2:* |  |  |  |  |  |  |  |  |  |  |  |  |  |
| *PCR: NP/CSF* | -/N,D, | -/N,D, | -/N,D, | N,D,/N,D, | N,D,/N,D, | N,D,/N,D, | N,D, | N,D, | N,D, | N,D, | N,D, | N,D, | N,D, |
| *IgG:* | -/- | -/- | -/- | -/- | -/- | -/- | N,D, | N,D, | N,D, | N,D, | N,D, | N,D, | N,D, |
| *Plasma/CSF* |  |  |  |  |  |  |  |  |  |  |  |  |  |
| *Used as control for:* |  |  |  |  |  |  |  |  |  |  |  |  |  |
| *Immunostaining* | YES | YES | YES | YES | YES | YES | NO | NO | NO | NO | NO | NO | NO |
| *IP-MS* | YES | YES | YES | YES | YES | YES | YES | YES | YES | YES | YES | YES | YES |
| *PhIP-Seq* | YES | YES | YES | YES | YES | YES | NO | YES | YES | NO | NO | NO | NO |

Table S3. Normalized gene counts for clusters in Figure 1.

Table S4. Differentially expressed genes in clusters described in Figure 1 between (a) COVID-19 patient CSF versus Control patient CSF. (b) COVID-19 patient PBMC versus Control patient PBMC. (c) COVID-19 patient CSF versus COVID-19 patient PBMC

Table S5. Normalized gene counts for T cell clusters in Figure 2.

Table S6. Differentially expressed genes between genes in T cell clusters described in Figure 2 (a) COVID-19 patient CSF versus Control patient CSF. (b) COVID-19 patient PBMC versus Control patient PBMC. (c) COVID-19 patient CSF versus COVID-19 patient PBMC

Table S7 Normalized gene counts for B cell clusters in Figure 3.

Data file S1. COVID-19 Case Histories

## Acknowledgements

We thank James A. Wells, James R. Byrnes and Jayant V. Rajan for their contribution of reagents to assist with the serologic assays. We are grateful to the participants who volunteered to be a part of this study, Patrick Wong and Orr-el Weizman for helpful discussion, Yale environmental health and safety team, and the Yale Center for Genome Analysis. **Funding:** This work was supported by NIH, K23MH118999 (SFF), F30CA239444 (ES), F30CA250249 (RDC), R01AI157488 (SFF and AI), R01AI104739 (SHK), U19AI089992 (SHK, AI), T32GM136651 (ES, RDC, RJ), K08NS096117 (MRW), R01MH122471 (JLD, SJP, MRW), R21MH118109 (SS), George Mason University (AI), Chan Zuckerberg Biohub (JLD), Brain Research Foundation (SJP), Program for Breakthroughs in Biomedical Research (SJP, MRW), Hanna H. Gray Fellowship, Howard Hughes Medical Institute (CMB), President’s Postdoctoral Fellowship Program, the University of California (CMB), John A. Watson Scholar Program, the University of California, San Francisco (CMB), the Beatrice Kleinberg Neuwirth Fund (AIK), and Fast Grant funding support from the Emergent Ventures at the Mercatus Center. LC-MS was supported by the NIH shared instrumentation grant S10OD021801.

## Author contributions

ES devised and executed single cell RNA sequencing and mouse SARS-CoV-2 experiments, analysis and interpretation of data resulting from said assays, and drafted the manuscript. CMB designed and assisted with anatomic immunostaining, immunoprecipitation mass spectrometry (IP-MS), phage display immunoprecipitation sequencing (PhIP-Seq), analysis and interpretation of data resulting from said assays, and wrote and edited the manuscript. RDC, RJ, and SHK assisted in analysis of single cell RNA sequencing. CRZ designed and performed experiments with SARS-CoV-2 Luminex assay. ACM, JC, CL, JK, HW, TM, BGI, and JO assisted in human subject recruitment and sample preparation. CBFV and NG contributed to SARS-CoV-2 RT-PCR and edited the manuscript. LM interpreted clinical data and edited the manuscript. SS and AIK provided human samples, edited the manuscript, and provided useful discussion. FL, YD, and AR assisted in SARS-CoV-2 ELISA and edited the manuscript. NTR provided bioinformatics analyses and edited the manuscript. TTN assisted with performing anatomic immunostaining, cloning of overexpression plasmids, overexpression cell-based assays, and microscopy, and interpretation of resulting data RPL assisted with performing immunoprecipitation mass spectrometry, SARS-CoV-2 Luminex assay, and cloning of monoclonal antibodies, and assisted with anatomic immunostaining. RD assisted with development of the PhIP-seq bioinformatic pipeline and PhIP-seq data presentation. IAH and BA assisted with performing PhIP-seq. BSP and MS assisted with IP-MS design, data acquisition, and analysis. JAG and TL recruited pre-pandemic human subjects. JLD assisted with development of the PhIP-seq platform, edited the manuscript, and provided useful discussion. AI assisted in data analysis, edited the manuscript, and provided useful discussion. SFF, SJP, and MRW conceived and supervised the project and wrote and edited the manuscript.

## Data availability

Gene expression and repertoire data in the study will be available at the time of publication in the NCBI repository SRAxxxx

